# *Arabidopsis* Myosins XI Are Involved in Exocytosis of Cellulose Synthase Complexes

**DOI:** 10.1101/492629

**Authors:** Weiwei Zhang, Chao Cai, Christopher J. Staiger

## Abstract

In plants, cellulose is synthesized at the cell surface by plasma membrane (PM)-localized cellulose synthase (CESA) complexes (CSCs). The molecular and cellular mechanisms that underpin delivery of CSCs to the PM, however, are poorly understood. Cortical microtubules have been shown to interact with CESA-containing compartments and mark the site for CSC delivery, but are not required for the delivery itself. Here, we demonstrate that myosin XI and the actin cytoskeleton mediate CSC delivery to the PM by coordinating the exocytosis of CESA-containing compartments. Measurement of cellulose content indicated that cellulose biosynthesis was significantly reduced in a myosin *xik xi1 xi2* triple knockout (*xi3KO*) mutant. By combining genetic and pharmacological disruption of myosin activity with quantitative live-cell imaging, we observed decreased abundance of PM-localized CSCs and reduced delivery rate of CSCs in myosin-deficient cells. These phenotypes correlated with a significant increase in failed vesicle secretion events at the PM as well as an abnormal accumulation of CESA-containing compartments at the cell cortex. Through high spatiotemporal assays of cortical vesicle behavior, we identified defects in CSC vesicle tethering and fusion at the PM. Furthermore, disruption of myosin activity reduced the delivery of several other secretory markers to the PM and reduced constitutive and receptor-mediated endocytosis. These findings reveal a previously undescribed role for myosin in vesicle secretion and cellulose production at the cytoskeleton–PM–cell wall nexus.

## INTRODUCTION

Cellulose microfibrils are the major load-bearing component of the plant cell wall and play essential roles in plant growth and development (McFarlane et al., 2014; Wallace and Somerville, 2015). Cellulose is produced at the plasma membrane (PM) by multimeric cellulose synthase complexes (CSCs), or rosettes, consisting of multiple cellulose synthase (CESA) proteins (Delmer, 1999; Somerville, 2006). Both freeze-fracture studies and live-cell quantitative imaging indicate that CSCs are assembled in Golgi (Giddings et al., 1980; Haigler and Brown, 1986; Paredez et al., 2006). CSCs are also present in small cytoplasmic CESA compartments (SmaCCs; Gutierrez et al., 2009) or microtubule-associated transport vesicles (MASCs; Crowell et al., 2009), which are associated with CSC delivery, generated by endocytosis, or both. Understanding the intracellular trafficking and delivery of CSCs is of great importance as it determines the abundance of CSCs at the PM and consequently affects the amount of cellulose produced and assembled in the cell wall (Bashline et al., 2014; Wallace and Somerville, 2015).

The cytoskeleton is implicated as a central player that coordinates trafficking of CSCs. In addition to choreographing the trajectory of CSCs in the PM, cortical microtubules interact with MASCs through the linker protein CELLULOSE SYNTHASE INTERACTIVE 1 (CSI1) and mark the sites for insertion of newly delivered CSCs (Paredez et al., 2006; Gutierrez et al., 2009; Bringmann et al., 2012; Zhu et al., 2018). However, they do not influence the rate of CSC delivery or abundance of CSCs at the PM, and cellulose content is not altered after treatment with the microtubule-disrupting drug oryzalin (Paredez et al., 2006; Gutierrez et al., 2009; Sampathkumar et al., 2013). In contrast, the actin cytoskeleton has recently been shown to participate in the delivery and endocytosis of CSCs, thereby affecting the amount of cellulose produced. SmaCCs are observed along subcortical actin filaments and translocate in an actin-dependent fashion (Sampathkumar et al., 2013). Genetic disruption of actin cytoskeleton organization in the *act2 act7* mutant or pharmacological perturbation with the actin polymerization inhibitor latrunculin B (LatB) leads to significant inhibition of the rate of delivery of CSCs to the PM and a marked reduction in overall cellulose content (Sampathkumar et al., 2013). In spite of these intriguing results, the molecular and cellular mechanisms that underpin a role for actin in vesicle delivery and CSC membrane dynamics remain unresolved.

In plant cells, a highly dynamic cortical actin network comprising single filaments and actin filament bundles is coordinated by a plethora of conserved and novel actin-binding proteins (Li et al., 2015). Myosins are molecular motors that transport diverse cargo along actin filaments and, in plants, are grouped into class XI and class VIII subfamilies (Vale and Milligan, 2000; Reddy and Day, 2001). In *Arabidopsis thaliana*, class XI contains 13 isoforms that share functional redundancy (Peremyslov et al., 2008). By analysis of *Arabidopsis* mutants with two, three or four of the most highly expressed *myosin XI* genes knocked out, these motors are implicated in regulating actin organization and dynamics and are key contributors to organelle and vesicle motility (Prokhnevsky et al., 2008; Peremyslov et al., 2010; Ueda et al., 2010; Peremyslov et al., 2012; Madison and Nebenführ, 2013; Cai et al., 2014). The velocity of myosin translocation along actin filaments, and therefore the rate of cytoplasmic streaming, also correlates with rates of cell expansion and plant growth (Tominaga et al., 2013). When chimeric myosins with faster or slower motor domains compared to endogenous myosins XI are expressed in transgenic *Arabidopsis* plants, the fast myosin supports larger epidermal cells and larger plants, whereas slow myosin leads to shorter cells and smaller plants (Tominaga et al., 2013). At present, it is not known how the velocity of chimeric myosins impact on axial cell expansion or whether this involves differences in cell wall composition and assembly.

In animal and yeast cells, actin and myosin are shown to participate in exocytosis and membrane fusion. For example, actin remodeling and/or polymerization correlates with increased vesicle secretion (Gutiérrez, 2012). In some systems, actin coats the secretory compartments and, together with myosin II, stabilizes the fusion pore or regulates the rate of fusion pore expansion (Nightingale et al., 2012). Furthermore, myosin V motors interact directly with Rab GTPases and the exocyst complex to facilitate secretory vesicle trafficking (Jin et al., 2011; Donovan and Bretscher, 2012). In plant cells, the function of actin and myosin in these processes has not been established. *Arabidopsis* Myosin XIK mediates the motility of several secretory vesicle markers, including the Rab GTPase RabA4b, the secretory carrier membrane 2 (SCAMP2) and the syntaxin of plants 41 (SYP41), and a functional XIK-YFP partially colocalizes with RabA4b and SCAMP2 compartments (Avisar et al., 2012; Peremyslov et al., 2012; Park and Nebenführ, 2013). A proteomics study in *Arabidopsis* identified both CESA and myosin XI in compartments enriched for the RabD2a/ARA5 marker, a Rab GTPase that decorates Golgi, TGN/EE and post-Golgi vesicles (Heard et al., 2015). However, other than vesicle motility, it is not known whether myosin XI also regulates secretion or fusion to the PM.

In this study, we combined genetic and pharmacological approaches with quantitative live-cell imaging to investigate the roles of myosin in CESA trafficking and CSC secretion. We demonstrated that cortical actin and myosin play a prominent role in vesicle exocytosis and are responsible for trafficking of CESA-containing compartments and tethering or insertion of CSCs at the PM.

## RESULTS

### Cellulose Content Is Reduced in the *myosin xi3KO* Mutant

An *Arabidopsis myosin xi1 xi2 xik* triple knockout mutant exhibits an overall dwarf plant phenotype with shorter cell lengths in both dark-grown hypocotyls and light-grown roots, resembling features that are typical of cellulose-deficient mutants and mimicking chemical inhibition of cellulose synthesis (Fagard et al., 2000; Peremyslov et al., 2010; Cai et al., 2014; Bashline et al., 2015). Although cellulose production involves intracellular trafficking and exocytosis of CSCs (Zhu et al., 2018), indications that myosins XI could be responsible are limited. To test whether myosin participates in cellulose production, we measured cellulose content in the previously characterized triple knockout mutant, hereafter referred to as *xi3KO* (Peremyslov et al., 2010). Cellulose content in alcohol-insoluble cell wall fractions of seedlings was determined after hydrolysis of all non-cellulosic material in acetic-nitric (AN) reagent (Updegraff, 1969); alternatively, hydrolysis in 2M trifluoroacetic acid (TFA) yields insoluble crystalline cellulose equivalent in amount to the AN reagent, but allows recovery of non-cellulosic monosaccharides including glucose from amorphous cellulose (Gibeaut and Carpita, 1991). The *xi3KO* mutant had significantly reduced total and crystalline cellulose compared to the wild-type control (Fig. 1). Similarly, wild-type seedlings grown on medium containing LatB showed a decrease in cellulose content (Fig. 1) and was consistent with a previous study (Sampathkumar et al., 2013). Further, seedlings grown on medium containing the plant myosin inhibitor 2,3-butanedione monoxime (BDM) also had significantly reduced cellulose (Supplemental Fig. S1).

**Figure 1.**
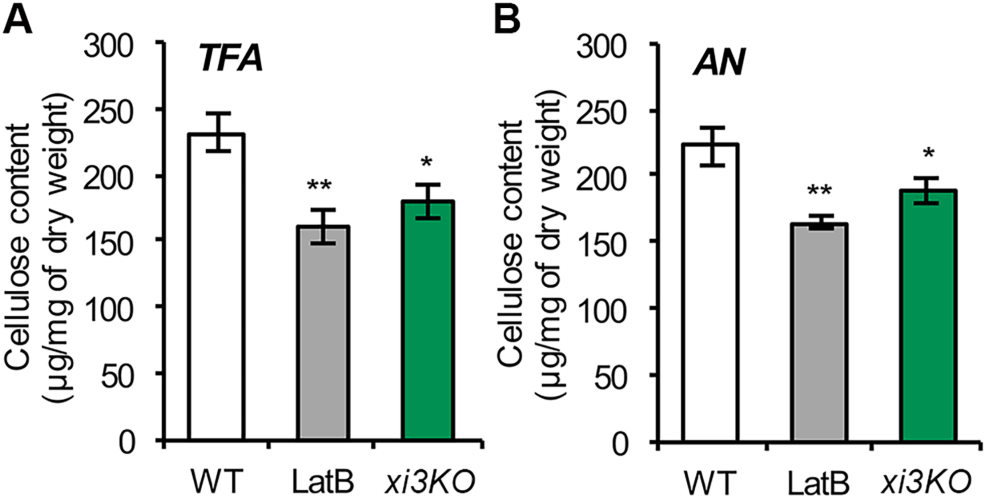
Cellulose content is reduced in the *myosin xi3KO* mutant. Ethanol-insoluble cell wall material (CWM) was prepared from 5-d-old etiolated hypocotyls of wild-type (WT) seedlings, the *myosin xi3KO* mutant, and WT growing on plates containing 100 nM latrunculin B (LatB). A and B, Cellulose content was measured as described in Methods. The non-cellulosic component of CWM was hydrolyzed with 2 M trifluoroacetic acid (TFA; A) for total cellulose determination, or with acetic nitric reagent (AN; B) for crystalline cellulose determination. Cellulose content was significantly reduced in LatB-treated and *xi3KO* mutant hypocotyls compared to WT. Values given are means ± SE (n = 4; Student’s *t* test, *P < 0.05, **P < 0.01).

We also analyzed cell wall monosaccharide composition to determine if non-cellulosic cell wall components were altered. No major differences in amounts of individual monosaccharides were detected in the cell wall fractions from seedlings of the *xi3KO* mutant or treated with LatB compared to those of wild type (Supplemental Table S1).

### Pentabromopseudilin is a Potent Myosin Inhibitor in Plant Cells

To investigate the role of myosin in cellulose deposition in detail, we used a pharmacological approach to acutely inhibit the activity of myosins. This further serves to bypass the potential genetic redundancy due to the presence of multiple myosins XI in *Arabidopsis*, as well as possible indirect effects resulting from long-term disruption of myosin activity in high-order mutants. BDM is the most commonly used plant myosin inhibitor; however, it requires a high concentration and its specificity has been questioned (McCurdy, 1999; Tominaga et al., 2000; Funaki et al., 2004). To search for additional plant myosin inhibitors, we screened chemical inhibitors of class V myosins, which are closely related to plant myosins VIII and XI (Foth et al., 2006). Pentabromopseudilin (PBP) and MyoVin-1 potently inhibit myosin V in animal and fungal cells (Fedorov et al., 2009; Islam et al., 2010).

To test whether these two inhibitors target plant myosins, we measured Golgi motility as a myosin XI-dependent process (Prokhnevsky et al., 2008; Peremyslov et al., 2010). Etiolated hypocotyl epidermal cells expressing yellow fluorescent protein (YFP) fused to mannosidase I, a Golgi resident enzyme that serves as a Golgi marker (Nebenführ et al., 1999), were imaged by time-lapse variable-angle epifluorescence microscopy (VAEM). After 15-min inhibitor treatment, fast directional and long-distance movement of Golgi was detected in mock and MyoVin-1 treated cells (Fig. 2, A and B; Supplemental Movie S1). In contrast, rapid and directional Golgi movement was markedly reduced in BDM-and PBP-treated cells, with most organelles moving slowly or wiggling in place (Fig. 2, A and B; Supplemental Movie S1). Quantitative analyses showed a significant reduction in average Golgi velocity after BDM and PBP treatment compared to control cells or those treated with MyoVin-1 (Fig. 2C). Plant Golgi display both fast directional and slow wiggling movement patterns (Akkerman et al., 2011). PBP appeared to inhibit both types of movement, whereas BDM mainly impacted the fast-moving population with a velocity greater than 1 µm s^−1^ (Supplemental Fig. S2). Moreover, we found that inhibition of Golgi motility by PBP was both time and dose dependent, and the drug could be washed out of cells to restore motility (Supplemental Fig. S3, A–C).

**Figure 2.**
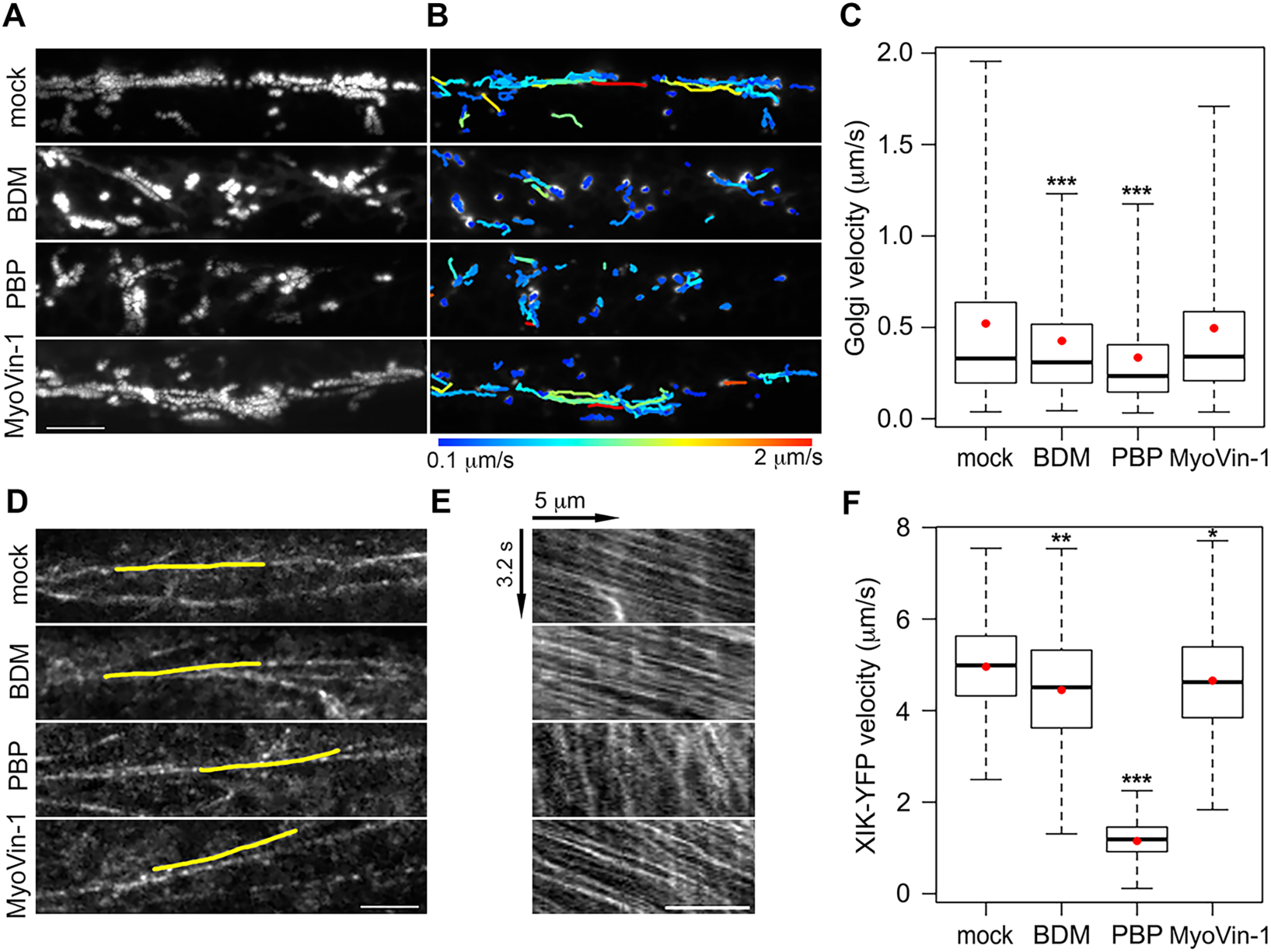
Myosin inhibitor treatments reduce motility in hypocotyl epidermal cells. A, Representative time projections show the trajectories of Golgi motility from apical epidermal cells of 3-d-old etiolated hypocotyls expressing YFP-Mannosidase I. Seedlings were treated for 15 min with mock (0.2% DMSO), 30 mM 2,3-butanedione monoxime (BDM), 10 µM pentabromopseudilin (PBP) or 20 µM MyoVin-1 prior to imaging. Time projections were generated with maximum intensity of 61 frames collected at 0.5-s intervals. Bar = 10 µm. B, Trajectories of Golgi motility detected with the ImageJ plugin “TrackMate” for images shown in (A). Heat map of trajectories indicates the average speed from 0.1 to 2 µm s–1. C, Box-whisker plots show average velocity of Golgi in inhibitor-treated epidermal cells. Red circles show the mean velocity. The rate of average Golgi velocity was significantly decreased after BDM and PBP treatments, whereas MyoVin-1 had little or no effect. Values given are means ± SE (n > 3000 trajectories from 10 hypocotyls per treatment; Student’s t test, ***P < 0.001). D, Representative single-frame images of the cortical cytoplasm in hypocotyl epidermal cells expressing Myosin XIK-YFP. Etiolated seedlings were treated with inhibitors for 15 min, followed by time-lapse imaging using an 80-ms interval for 41 frames. Bar = 5 µm. E, Kymographs of the yellow lines depicted in (D) show fast movement of the vesicles over a 3.2 s timespan. Bar = 5 µm. F, Quantification of XIK-YFP velocity from analysis of multiple kymographs. The motility of YFP-XIK was markedly decreased in PBP-treated cells and slightly reduced in BDM or MyoVin-1-treated cells compared to mock treatment. Values given are means ± SE (n > 500 tracks from 10 hypocotyls per treatment; Student’s *t* test, *P < 0.05, **P < 0.01, ***P < 0.001).

To test the effects of these inhibitors on myosin-based motility more directly, we took advantage of a functional, full-length Myosin XIK fused with YFP (XIK-YFP; Peremyslov et al., 2012). This reporter associates with small vesicle clusters that traffic along F-actin and appear as “beads-on-a-string” (Peremyslov et al., 2012). We measured velocities of these vesicles after inhibitor treatment using time-lapse images collected with spinning-disk confocal microscopy (SDCM). In mock-treated cells, the XIK-YFP vesicles moved at a mean speed of ∼5 µm s^−1^, whereas mean velocities decreased by nearly 80% in PBP-treated cells (Fig. 2, D–F; Supplemental Movie S2). In contrast, mean velocities decreased by only 10% upon BDM treatment, and less than 5% with MyoVin-1 treatment (Fig. 2, D–F; Supplemental Movie S2). The reason that BDM did not strongly inhibit XIK-YFP motility could be that it has a lower specificity in targeting the XIK isoform. Finally, PBP showed a dose-and time-dependent inhibition of XIK-YFP motility and the drug could be washed out of cells to restore motility (Supplemental Fig. S3, D–F).

Myosin XI in the green alga *Chara corallina* has the fastest velocity reported among all myosin motors (∼50 µm s^−1^) and drives rapid cytoplasmic streaming (Yamamoto et al., 1994; Shimmen and Yokota, 2004). PBP inhibited the velocity of cytoplasmic streaming in internodal cells of *C. corallina* by ∼75% after 15 min treatment with 1 µM PBP and the inhibition was both dose-and time-dependent (Supplemental Fig. S4). Taken together, these data demonstrate that PBP effectively inhibits myosin-dependent organelle motility and validates its use as a myosin inhibitor in plants. In contrast, MyoVin-1 did not inhibit any plant myosin-dependent process tested herein (Supplemental Table S2) and will not be used further.

### Abundance of Cellulose Synthase Complexes in the Plasma Membrane Is Reduced in the *xi3KO* Mutant

One explanation for the reduced cellulose content in *xi3KO* could be that the number of cellulose synthase complexes (CSCs) in the plasma membrane (PM) is decreased. To address this possibility, a line expressing YFP-CESA6 in the *prc1-1* homozygous mutant background (Paredez et al., 2006) was crossed with the *xi3KO* mutant. We recovered *myosin xi* triple homozygous knockouts as well as wild-type siblings expressing YFP-CESA6 in the presence of *prc1-1*. When examined by SDCM, the density of CSCs at the PM in the *xi3KO* mutant was reduced by greater than 30% compared to wild-type siblings (Fig. 3, A–C). We also performed short-term treatment of wild-type siblings with BDM and PBP to test whether acute inhibition of myosin activity could recapitulate the reduction of CSC density in the *xi3KO* mutant. After BDM and PBP treatment for 15 min, the density of PM-localized CSCs dropped significantly by 14% and 25%, respectively, compared to that of non-treated cells (Fig. 3, A–C). In contrast, treatment with LatB at a high concentration (10 µM) had no effect on CSC density (Fig. 3, A–C); the latter result was consistent with a previous report (Sampathkumar et al., 2013). The motility of CSCs in the plane of PM was also reduced in the *xi3KO* mutant and BDM-and PBP-treated cells, but not in LatB-treated cells (Fig. 3, D–F).

**Figure 3.**
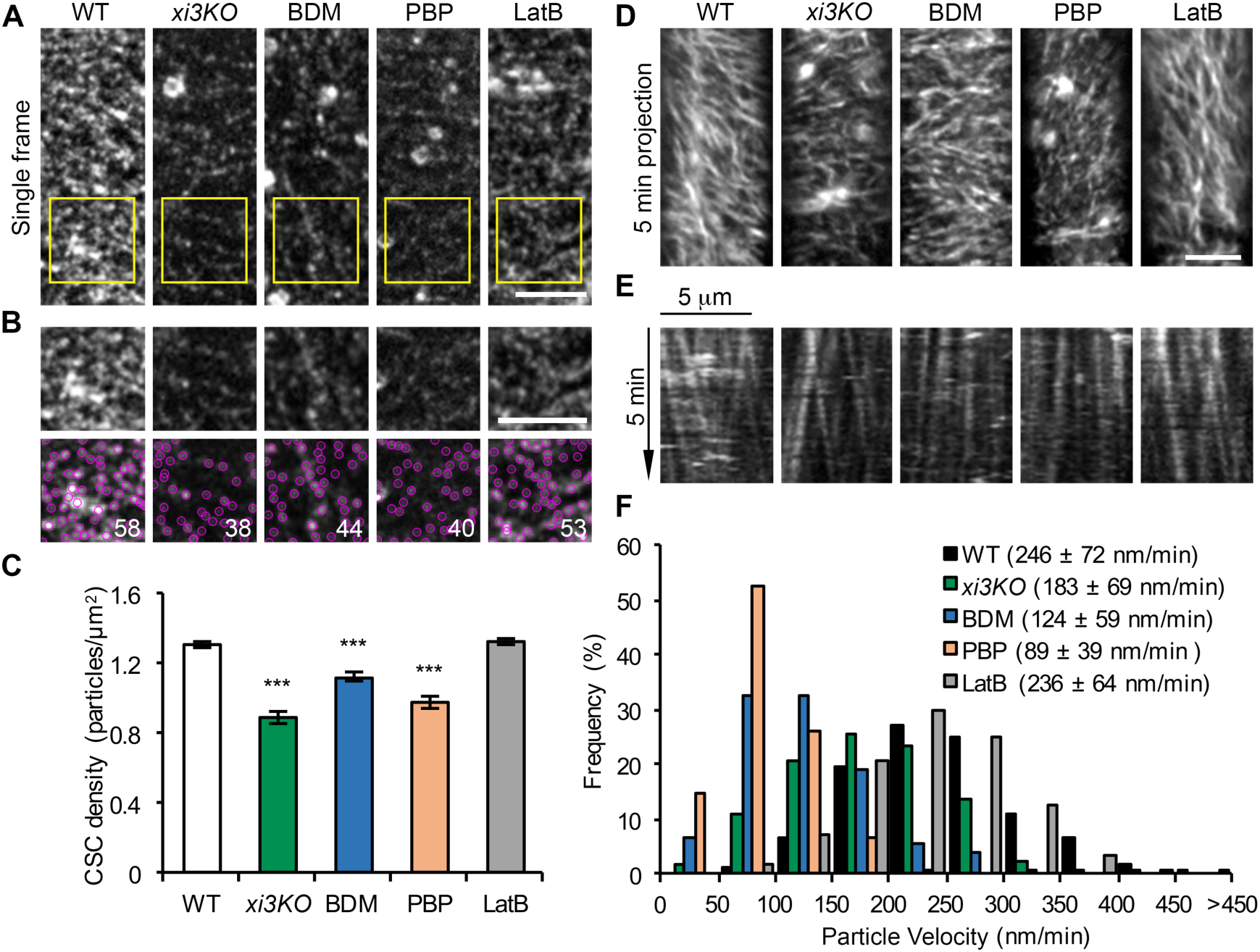
The density and motility of CSC particles at the plasma membrane are reduced in myosin-deficient cells. Etiolated hypocotyl epidermal cells of *xi3KO* or wild-type (WT) siblings expressing YFP-CESA6 were imaged with spinning disk confocal microscopy. Seedlings were pre-treated with mock (0.2% DMSO), 30 mM BDM, 10 µM PBP or 10 µM LatB for 15 min. A, Representative single-frame images show the distribution of CSCs at the plasma membrane (PM). Bar = 5 µm. B, Selected regions (yellow box) in (A) were magnified and CSC particles detected (marked in magenta). The number of particles in each region is shown in white. Bar = 5 µm. C, Quantitative analysis shows that the density of CSC at the PM was significantly reduced in *xi3KO* or after BDM and PBP treatment, but not in LatB-treated cells. Values given are means ± SE (n ≥ 35 cells from 10 hypocotyls per treatment; more than 12,000 particles were measured from total areas of > 9,400 µm2 in WT, *xi3KO*, BDM-, PBP-and LatB-treated cells. Student’s *t* test, ***P < 0.001). D, Representative time projections show the trajectories of CSCs at the PM. Time projections were generated by average intensity with 61 frames collected at 5-s intervals. Bar = 5 µm. E, Representative kymographs show the movement of CSCs during 5 min interval. F, Quantitative analysis shows the distribution and average velocities of CSCs. The average CSC velocities in *xi3KO*, BDM-and PBP-treated cells were greatly reduced compared to WT or LatB-treated cells. Values given are means ± SD (n > 500 CSC trajectories per treatment).

The pattern of CESA distribution at the PM correlates with the position of Golgi in the cortical cytoplasm and disruption of the actin cytoskeleton reportedly causes clustering of CESA-containing Golgi bodies, and consequently, uneven distribution of CSCs at the PM (Gutierrez et al., 2009; Sampathkumar et al., 2013). We observed similar patterns in LatB-treated cells, whereas in *xi3KO* or cells treated with BDM or PBP, apparent Golgi clustering or patchy distribution of CSCs at the PM was not detected frequently (Supplemental Fig. S5). In support of these observations, we quantified the percentage of single Golgi bodies in the cell cortex as an indicator of the level of Golgi clustering. In LatB-treated cells, there was a significantly lower proportion of single Golgi compared with wild type, confirming a high extent of Golgi clustering, whereas in *xi3KO*, BDM-and PBP-treated cells the proportion of single Golgi increased significantly (Supplemental Fig. S5). This indicates that, in contrast to actin, disruption of myosin does not cause clustering of cortical Golgi and the decrease of CSC density at the PM in myosin-deficient cells is unlikely a result of disruption of Golgi distribution.

### Rate of Delivery of CSCs to the PM Is Determined by Myosin

The decreased density of CSCs in myosin-deficient cells indicates that myosin may be required for the delivery of CSCs to the PM. Fluorescence recovery after photobleaching (FRAP) experiments have been developed to analyze the rate of delivery of CSCs to the PM (Gutierrez et al., 2009; Bashline et al., 2013; Sampathkumar et al., 2013; Luo et al., 2015). To test the role of myosin, we performed FRAP experiments on the *xi3KO* mutant expressing YFP-CESA6.

Briefly, a small region was irradiated to minimize the bleaching of the cytoplasmic mobile CESA pool, and a central sub-region was selected to avoid error introduced by migration of CSCs into the measured region (Fig. 4A). For inhibitor studies, to minimize the duration of treatment, we did not pretreat hypocotyls but instead mounted directly in drug and immediately began time-lapse imaging. It has been reported that the rate of CSC delivery correlates with the abundance of underlying CESA-containing Golgi and choosing a bleached region with few Golgi will affect the apparent delivery frequency (Gutierrez et al., 2009; Sampathkumar et al., 2013). To reduce the chances of biasing our analysis toward regions of PM that were deficient in CSC delivery due to a lack of nearby Golgi, we avoided regions that had few CSC particles or few Golgi. This was confirmed by analyzing the density of CSC in regions selected for photobleaching, and no significant difference was detected between any genotype or treatment (Supplemental Fig. S6).

**Figure 4.**
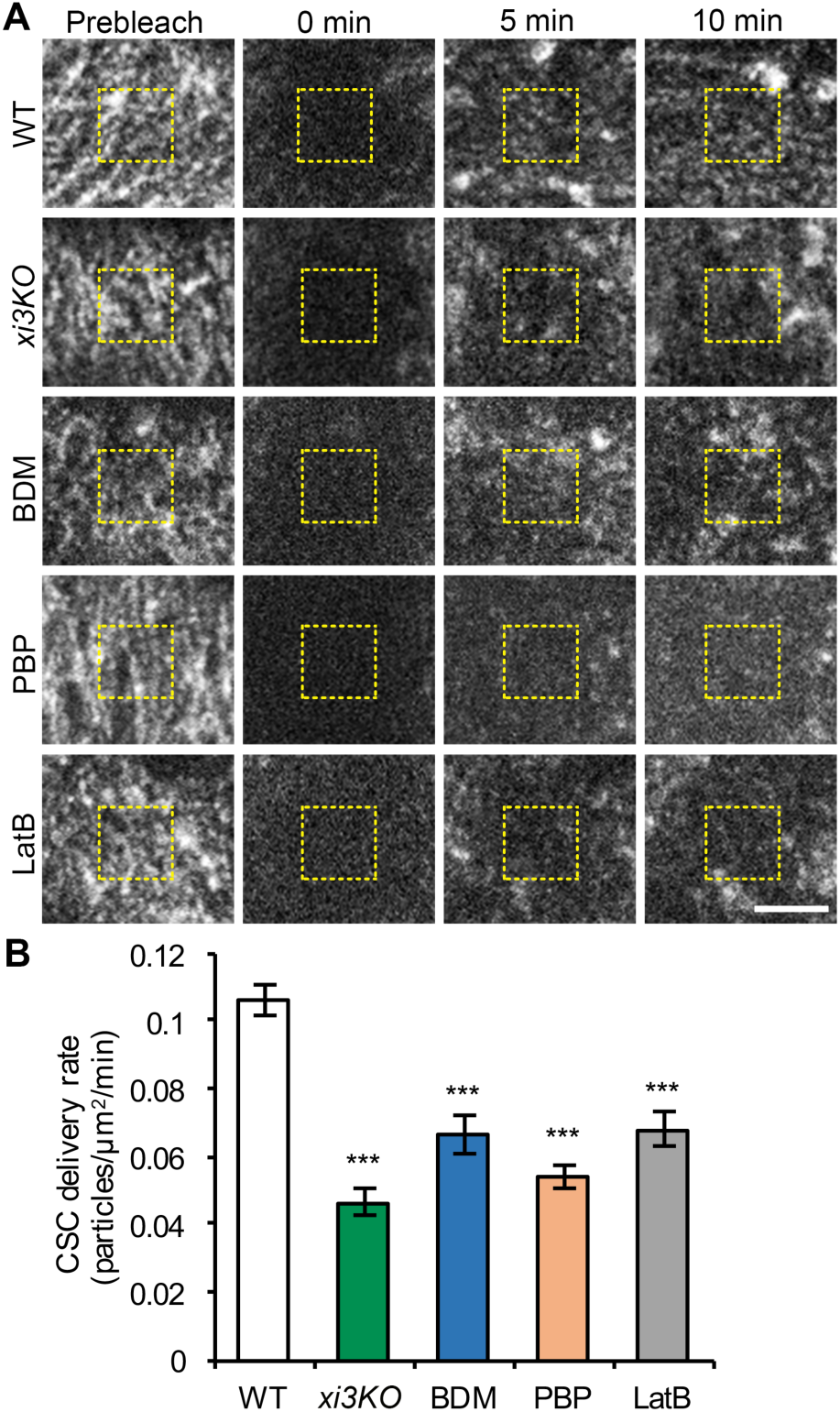
Delivery of CSCs to the PM is reduced in myosin-deficient cells. A, Representative single-frame images of PM-localized CSC particle recovery after photobleaching. Seedlings of WT or *xi3KO* were mounted in mock (0.2% DMSO), 30 mM BDM, 10 µM PBP or 10 µM LatB just prior to imaging. A region of interest at the PM was photobleached and the number of newly-delivered CSCs was counted in a subarea within the region (yellow dashed box). Bar = 5 µm. B, The rate of delivery of CSCs to the PM was calculated from the total number of newly-delivered CSCs during the initial 5 min of recovery divided by the measured area and time. The CSC delivery was significantly inhibited in *xi3KO*, BDM-, PBP-and LatB-treated cells compared with mock-treated WT sibling. Values given are means ± SE (n ≥ 11 cells per genotype or treatment, Student’s *t* test, ***P < 0.001).

After photobleaching, we observed gradual recovery of CSCs in all genotypes or treatments; however, the rate of recovery in *xi3KO*, BDM-, PBP-and LatB-treated cells was reduced compared with WT (Fig. 4A). We quantified the newly delivered CSC particles in the bleached region according to published methods and only *de novo* appearance of particles that exhibited a static phase followed by steady migration were counted as new delivery events (Gutierrez et al., 2009). Mock-treated wild-type cells had a delivery rate of 0.11 ± 0.02 particles µm^−2^ min^−1^ (Fig. 4B). In contrast, the CSC delivery rate in the *xi3KO* mutant decreased by greater than 50% (Fig. 4B). In BDM-, PBP-and LatB-treated cells, the delivery rates were reduced by 37%, 48%, and 36%, respectively (Fig. 4B). The reduced rate of delivery of CSCs in both *xi3KO* and myosin inhibitor-treated cells indicates that myosin is involved in delivery or insertion of CESA at the PM.

To verify that cortical microtubules do not affect that rate of delivery of CSC (Gutierrez et al., 2009), we pre-treated seedlings with oryzalin for 2 h which destabilized the majority of microtubules (Supplemental Fig. S7A). FRAP analysis of oryzalin-treated cells showed that the CSC delivery rate was not significantly different compared to control cells (Supplemental Fig. S7, B–D).

We also tested whether the actin-myosin transport network mediates delivery of general cargo proteins to the PM, in addition to CESA. We obtained two well-characterized protein markers, the auxin efflux carrier PINFORMED2 (PIN2) and the brassinosteroid receptor BRASSINOSTEROID INSENSITIVE1 (BRI1), which are actively delivered to the PM through *de novo* secretion or endomembrane recycling (Geldner et al., 2007; Luschnig and Vert, 2014). Treatment of roots expressing PIN2-GFP or BRI1-GFP with BDM, PBP or LatB for 2 h revealed significantly reduced fluorescence intensity at the PM, compared with that in mock-treated cells (Supplemental Fig. S8, A, B, D and E). The decreased PM fluorescence intensity correlated with an increased number of intracellular compartments (Supplemental Fig. S8, A and B), suggesting that the delivery of PIN2-GFP and BRI1-GFP to the PM was inhibited and the proteins were retained intracellularly after myosin-or actin-inhibitor treatment. In addition, treatment of cells expressing the secretory marker secGFP with BDM and PBP caused a significant retention of secGFP signal inside the cell, compared with complete secretion out of the cell in mock-treated cells (Supplemental Fig. S8, C and F). Collectively, our results indicate that myosin activity is required for general protein secretion, including delivery of CSCs to the PM.

### CESA Compartments Accumulate in the Cortical Cytoplasm After Myosin Disruption

Small cytoplasmic CESA-containing compartments or vesicles observed at the cell cortex are likely to be associated with delivery of CSCs to the PM through a poorly-defined mechanism (Crowell et al., 2009; Gutierrez et al., 2009; Wallace and Somerville, 2015). Previous work showed association of CESA compartments with actin filaments in the subcortical cytoplasm (Sampathkumar et al., 2013). We tested whether myosin XI functions in trafficking of CESA compartments to the PM for CSC delivery. When we segmented optical sections into cortical (0 to 0.4 µm below the PM) and subcortical (0.6 to 1 µm below the PM) cytoplasm (Sampathkumar et al., 2013), we found that the abundance of cytoplasmic CESA compartments in the cortical cytoplasm increased more than two-fold over wild-type cells in the *xi3KO* mutant or after acute treatment with BDM or PBP (Fig. 5). Treatment with LatB also increased the number of cortical CESA compartments to a similar level (Fig. 5). In contrast, the number of subcortical CESA compartments was either unchanged or was modestly increased upon drug treatment (Fig. 5).

**Figure 5.**
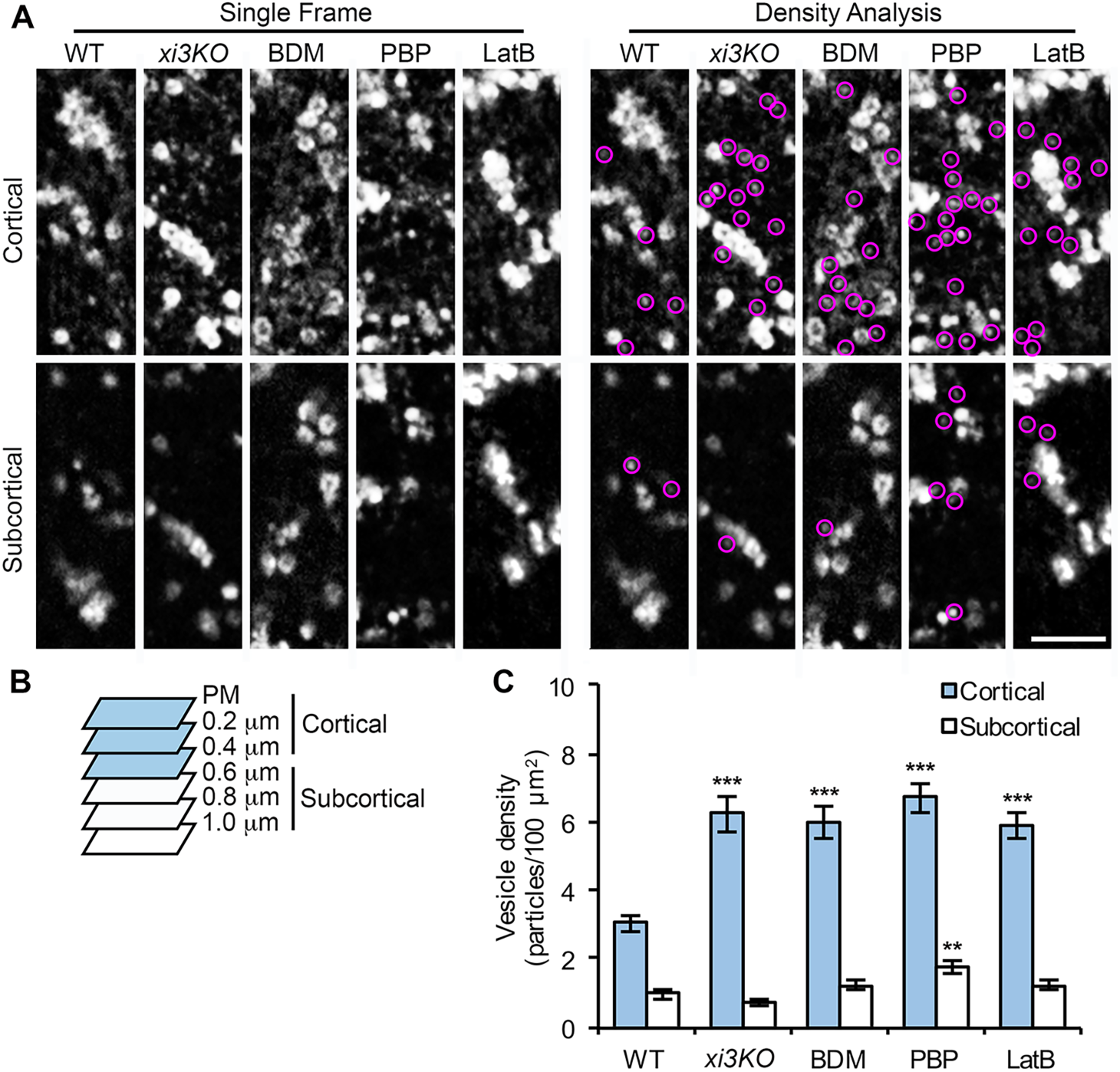
The density of small CESA compartments is increased in the cell cortex following myosin inhibition. A, Representative single frames taken at cortical and subcortical focal planes in hypocotyl epidermal cells. Seedlings of *xi3KO* or WT siblings expressing YFP-CESA6 were treated with mock (0.2% DMSO), 30 mM BDM, 10 µM PBP or 10 µM LatB for 15 min prior to imaging. The cytoplasmic CESA compartments are highlighted with magenta circles. Bar = 5 µm. B, Diagram shows that optical sections of 0.2-µm step size were taken beginning at the PM. Focal planes from the PM to 0.4 µm below the PM were defined as cortical cytoplasm and from 0.6 µm to 1.0 µm were defined as subcortical cytoplasm. C, Quantitative analysis of vesicle density shows that the number of CESA compartments was increased significantly in the cortical but not the subcortical cytoplasm in *xi3KO*, BDM-, PBP-and LatB-treated cells. Values given are means ± SE (n ≥ 20 cells from 10 seedlings for each genotype or treatment; a total of 333, 567, 790, 686 and 602 compartments were measured from total areas of 8344, 8418, 11229, 8321 and 8041 µm2 in WT, *xi3KO*, BDM-, PBP-and LatB-treated cells, respectively. Student’s *t* test, **P < 0.01, ***P < 0.001).

An increased abundance of cortical MASCs has been observed in cells treated with cellulose synthesis inhibitors (e.g. isoxaben) or osmotic stress (Crowell et al., 2009; Gutierrez et al., 2009). Under these conditions, CSCs were mostly depleted from the PM and internalized into MASCs which were tethered to cortical microtubules and displayed stationary or steady microtubule tip-tracking motility with an average velocity of ∼4 µm min^−1^ (Crowell et al., 2009; Gutierrez et al., 2009; Sampathkumar et al., 2013; Lei et al., 2015). In contrast, the accumulated CESA compartments we observed in myosin-and actin-deficient cells exhibited high dynamicity (Supplemental Movie S3). Most of the compartments displayed erratic or diffusive movement with mean velocities greater than 7 µm min^−1^ in all genotypes or inhibitor treatments, and we did not observe the stationary or microtubule-dependent motility pattern of CESA compartments in the cell cortex (Supplemental Fig. S9, A and C; Supplemental Movie S3). The motility pattern of those compartments suggests that they were not the same population as the MASCs observed in cells treated with isoxaben or under stress. In addition, inhibition of myosin or actin only reduced the fast-moving population of CESA compartments (> 20 µm/min) in both cortical and subcortical cytoplasm, but not those that had a lower motility (Supplemental Fig. S9), suggesting that the fast movement of the vesicles is dependent on actin and myosin activity but the vesicles could still retain motility, possibly through free diffusion when actin or myosin activity is disrupted.

The increased number of cytoplasmic CESA compartments in myosin-and actin-deficient cells could result from exocytic vesicles whose secretion was disrupted, from increased internalization of CESAs through endocytosis, or both. To test whether endocytosis was altered in *xi3KO*, BDM-, PBP-and LatB-treated cells, we utilized the lipophilic dye FM4-64, a marker for constitutive endocytosis, for an internalization assay in light-grown hypocotyl epidermal cells. The number of FM4-64-labeled endosomes was significantly reduced in the mutant and in cells pre-treated with BDM, PBP or LatB for 30 min (Supplemental Fig. S10, A and C). These results were consistent with a previous study showing that endocytosis of FM4-64 was greatly inhibited in a myosin *xi1 xi2 xii xik* quadruple knockout mutant (Yang et al., 2014).

BDM has also been shown to inhibit the ligand-induced endocytosis of the PM defense receptor FLAGELLIN SENSING2 (FLS2; Beck et al., 2012). We quantified the number of internalized FLS2 endosomes by pre-treating an *Arabidopsis* FLS2-GFP transgenic line with BDM or PBP followed by elicitation with the flagellin-derived ligand, flg22. Both BDM and PBP treatment greatly reduced the number of internalized FLS2 endosomes compared with mock treatment (Supplemental Fig. S10, B and D). The internalization assays indicated that perturbation of myosin activity inhibited both constitutive and receptor-mediated endocytosis, and therefore, the cortical CESA compartments that accumulated in myosin-and actin-deficient cells were more likely to be vesicles that were unable to deliver CESAs to the PM rather than newly internalized endosomes. These data support a role of cortical actin and myosin in mediating exocytic trafficking of CESA compartments for CSC delivery.

Because disruption of myosin activity alters actin organization and dynamics (Park and Nebenführ, 2013; Cai et al., 2014), we tested whether the accumulation of cortical CESA compartments in *xi3KO* and myosin inhibitor-treated cells resulted from perturbation of the actin cytoskeleton. We analyzed the density and extent of bundling of actin arrays in the cortical and subcortical cytoplasm, respectively. In wild-type cells, the density of actin filaments was significantly higher and less bundled in the cortex compared with that in the subcortex (Supplemental Fig. S11). In *xi3KO*, BDM and PBP-treated cells, a similar overall pattern of cortical and subcortical actin architecture was observed, although the average density was decreased and extent of bundling was increased in the mutant and PBP-treated cells. In contrast, cells treated with BDM had the opposite phenotype with higher filament densities in both the cortex and subcortex and reduced extent of bundling, compared with wild-type cells (Supplemental Fig. S11, Supplemental Table S2). These data indicate that the preferential accumulation of CESA compartments in the cortex in *xi3KO* or myosin inhibitor-treated cells is unlikely due to a specific alteration of actin architecture.

The SNARE protein Syntaxin of Plants 61 (SYP61) is a *trans*-Golgi network (TGN) marker and SYP61 compartments have been implicated in exocytic trafficking and transport of various cargoes, including CESA (Drakakaki et al., 2012). After treatment of etiolated hypocotyls expressing SYP61-CFP with BDM, PBP and LatB, we detected a significant increase in the number of small SYP61 vesicles in the cell cortex, but not in the subcortex, compared with control cells (Supplemental Fig. S12). These results reveal that actin and myosin are also involved in the trafficking of SYP61 vesicles and may play a role in general exocytic membrane trafficking.

Previous studies have shown that a population of CESA compartments co-localize with SYP61 (Gutierrez et al., 2009). To test whether the accumulated cortical CESA and SYP61 compartments in myosin-and actin-deficient cells are identical, we generated a line co-expressing YFP-CESA6 and SYP61-CFP and performed a co-localization assay. After BDM, PBP and LatB treatment, the abundance of both YFP-only and CFP-only compartments increased significantly, whereas the population with overlapping signals remained constant (Fig. 6, A and B). In mock-treated cells, 33% of the cortical vesicles had both YFP and CFP signals, however, after inhibitor treatments, the percentage of vesicles with overlapping signals was reduced to 20% or less (Fig. 6C). These results demonstrate that cortical CESA and SYP61 compartments that accumulate following myosin inhibition are distinct populations that do not significantly overlap.

**Figure 6.**
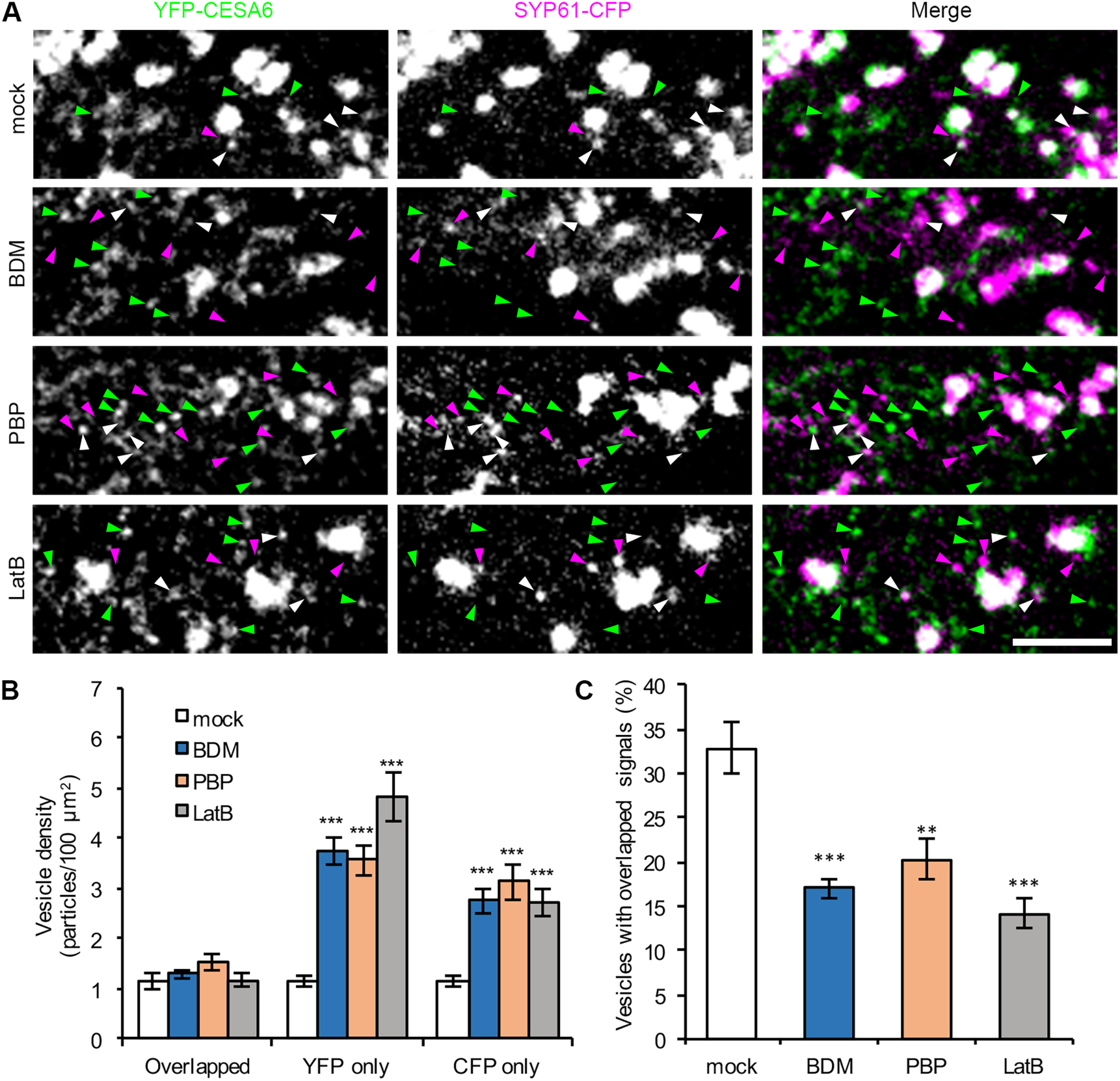
Myosin inhibitors cause an accumulation of distinct CESA6 and SYP61 compartments in the cortical cytoplasm. A, Representative single optical sections taken from the cortical cytoplasm in etiolated hypocotyl epidermal cells co-expressing YFP-CESA6 and SYP61-CFP. Seedlings were treated with inhibitors for 15 min prior to imaging. Vesicles labeled with YFP only, CFP only, or with both markers, are indicated by green, magenta and white arrowheads, respectively. Bar = 5 µm. B, Quantitative analysis of vesicle density shows that the number of YFP-only and CFP-only compartments was significantly increased after BDM, PBP and LatB treatment; in contrast, the number of compartments with overlapping signals did not show major differences compared with mock-treated cells. C, The percentage of vesicles with overlapping signals decreased significantly in BDM-, PBP-and LatB-treated cells. Values given are means ± SE (n ≥ 20 cells from 6 seedlings for each treatment; a total of 263, 586, 529 and 578 compartments were measured from total areas of 7804, 7649, 6339 and 6880 µm2 in mock, BDM-, PBP-and LatB-treated cells, respectively. Student’s *t* test, **P < 0.01, ***P < 0.001).

### Myosin Disruption Results in CSC Secretion Failure at the PM

One possibility for the abnormal accumulation of motile CESA compartments near the PM in myosin-and actin-deficient cells is that their secretion was inhibited. To test whether myosin plays a role in vesicle exocytosis, we observed individual CSC insertion events at the PM in epidermal cells from the middle region of the hypocotyl, where CSC particles are less dense and visualization of individual particle insertion was possible (Crowell et al., 2009). For the CSC insertion assay, seedlings were mounted directly in mock or inhibitor solutions to minimize inhibitor treatment time and time-lapse series were acquired at 3-s intervals for 12 min. Similar to a previous report (Gutierrez et al., 2009), we observed that a newly arrived CSC particle typically displayed three phases of movement: 1) a short erratic phase with rapid and local movements near the PM, 2) a static or pausing phase for 1–2 min in a fixed location, and 3) a steady migration phase as active cellulose-producing complexes translocate in the plane of the PM (Fig. 7A, Supplemental Movie S4). We also observed that most of the CSC insertion sites were coincident with cortical microtubules (Supplemental Fig. S13, Supplemental Movie S5), which was consistent with a previous report (Gutierrez et al., 2009).

**Figure 7.**
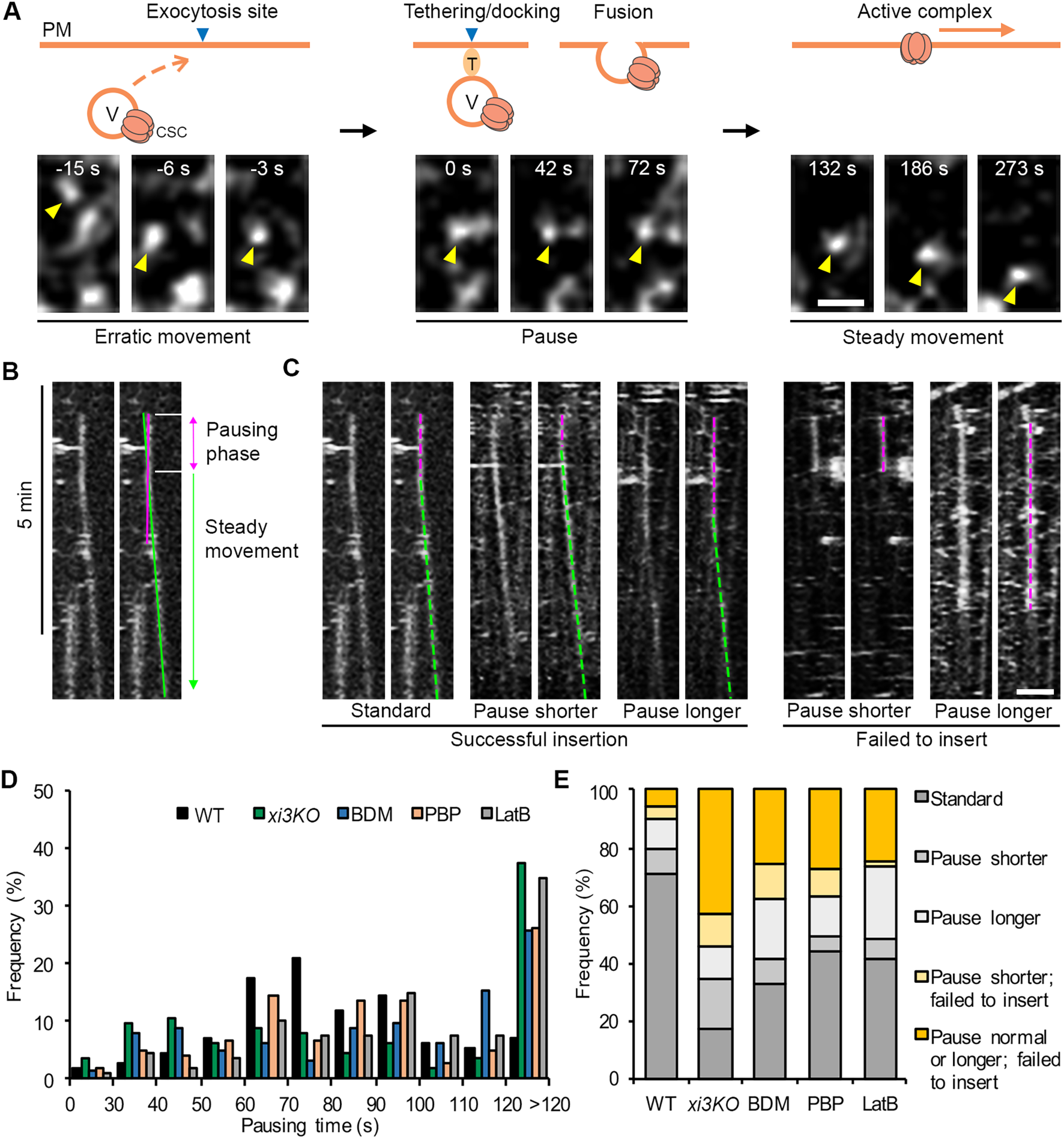
Inhibition of myosin causes CSC insertion failures at the PM. A, Representative images show a typical CSC insertion event at the PM. A CESA particle (yellow arrowhead) arriving in the cortex initially undergoes erratic motility, which likely represents a delivery vesicle (V) that is transported to an exocytosis site. The particle then pauses (marked as 0 s) and exhibits a static phase for ∼80 s in a fixed position, which likely corresponds to tethering, docking and fusion of the delivery compartment to the PM. After the CSC particle is inserted, it shows steady movement at the PM as an active complex. The full time series is shown in Supplemental Movie 4. T: tethering proteins. Bar = 1 µm. B, Representative kymograph of the CSC insertion event shown in (A) illustrates the pausing phase and the steady movement phase. The exact duration of the pausing phase was determined by fitting a straight line (green) along the moving trajectory and another line (magenta) along the pausing phase. The intersection of the two lines was defined as the end of the pausing phase. C, Representative kymographs show five categories of insertion events from left to right: standard insertion with normal pausing time; insertion that has a shorter pausing time; insertion that has a longer pausing time; a shorter pausing time and failure to insert; and, a longer pausing time and failure to insert. Pausing phases are marked with magenta dashed lines and steady movement phases are marked with green dashed lines. Bar = 2 µm. D, Distribution of particle pausing times. (n ≥ 11 cells from 7 seedlings for each genotype or treatment; a total of 114, 113, 124, 104 and 109 events were measured in WT, *xi3KO*, BDM-, PBP-and LatB-treated cells, respectively.) E, The proportion of five types of insertion events described in (C) in WT, *xi3KO*, BDM-, PBP-and LatB-treated cells. A shorter or longer pausing time was defined as the mean value (81 ± 27 s) of particle pausing time in WT minus (< 54 s) or plus one standard deviation (> 108 s), respectively.

The initial erratic phase of CSC particles is thought to represent the transient behavior of trafficking compartments before they dock and fuse to the destination PM to deliver CSCs (Gutierrez et al., 2009). In this study, we observed a total of 564 insertion events and 378 of them (67%) were associated with an erratically moving CESA compartment; these compartments often translocated in close association with Golgi, or were separate from a Golgi body, before they stabilized and paused at the PM (Fig. 7A; Supplemental Movies S4, S5, S6 and S7). The frequency of the erratic phase was similar to a previous report (37 out of 60; Gutierrez et al., 2009). In the rest of the delivery events, either pausing of a CESA-containing Golgi was observed to coincide with a new insertion or no obvious delivery compartments/organelles were observed. Our results confirm that the cytoplasmic CESA vesicles are the major delivery compartments for CSC localization in the PM.

The exocyst complex and an exocytosis-related protein PATROL1 were shown to colocalize with CSCs during the first few seconds of the pausing phase (Zhu et al., 2018), suggesting that pausing is a necessary step related to vesicle tethering and fusion for CSC delivery to the PM. Thus, measuring the pausing phase represents a way to assess the secretion process of CSCs. In this assay, only particles that showed *de novo* appearance at the PM plane followed by a static phase for more than 5 frames (> 15 s) were tracked and considered as insertion or attempted insertion events. By tracking the newly inserted particles over time, we found that the insertion events mainly fell into the following categories: (i) particles that paused for 1–2 min followed by steady movement as an active CSC complex (Fig. 7C, Supplemental Movie S4); (ii) particles that paused for a significantly shorter or longer time, followed by steady migration as an active CSC complex (Fig. 7C); and (iii) particles that underwent a pausing phase but then disappeared or rapidly moved away, therefore no CSC was inserted, representing a failed insertion/secretion event (Fig. 7C, Supplemental Movie S6, S7). We quantified each type of event and measured particle pausing time by analysis of kymographs (Fig. 7B). In wild type, the average CSC pausing time was 81 ± 27 s (n = 110 events), which was close to the previously reported value (62 ± 23 s, Gutierrez et al., 2009). Notably, particle pausing time was greatly altered in *xi3K*O or inhibitor-treated cells (Fig. 7D). To define aberrant pausing, we used the mean value of particle pausing time in wild type minus or plus one standard deviation; thus, pausing times < 54 s or > 108 s were considered as shorter or longer pausing events, respectively. We found that in *xi3K*O or inhibitor-treated cells, more than 30% of the insertions exhibited a longer pausing time (> 108 s); moreover, there was a slightly increased population that paused only for 30–50 s in *xi3KO*, BDM-and PBP-treated cells (Fig. 7D). Overall, the percentage of failed insertion events increased by 2–5 fold in the mutant or inhibitor-treated cells compared with wild type (Fig. 7E). In wild type, only 10% of the observed events failed to deliver a functional CSC (11 of 114 events), however, in *xi3KO,* the proportion increased to 55% (62 of 113 events) and upon BDM, PBP and LatB treatment, it increased to 37% (46 of 124 events), 37% (38 of 104 events) and 26% (28 of 109 events), respectively (Fig. 7E). The abnormal CSC pausing time and increased insertion failures at the PM indicate that cortical actin and myosin activity are required for exocytosis of CESA compartments, probably through mediating vesicle tethering or fusion to the PM.

Cortical microtubules have been shown to define the CSC insertion sites and CESA compartments are tethered to microtubules through CSI1 for CSC delivery (Gutierrez et al., 2009; Bringmann et al., 2012; Zhu et al., 2018). Another report showed that the microtubule-localized insertion of CSCs is not affected in LatB-treated cells or in the *act2 act7* double mutant (Sampathkumar et al., 2013). We tested whether disruption of myosin might affect the preferential insertion of CSCs next to microtubules, and our results showed that after treatment with BDM, PBP or LatB, most of the CSC insertion events (∼85%) observed were still coincident with microtubules, at a frequency similar to that in untreated cells (Supplemental Fig. S13). Therefore, myosin activity is not necessary for CSCs to reach the cortical microtubule sites but is required for the later stages of vesicle tethering and fusion to the PM.

## DISCUSSION

In plants, a cortical actin-myosin cytoskeletal network regulates cytoplasmic streaming, long-distance transport of diverse intracellular organelles and vesicles, and promotes cell expansion and growth. However, it remains unclear whether actin and myosin also direct vesicle exocytosis and protein secretion at the PM. In this study, by investigating the trafficking of CSC complexes, we demonstrate a previously undiscovered role for myosin XI in post-Golgi vesicle trafficking and cell wall synthesis. The large size of individual CSC particles as well as their organized spatial distribution and motility pattern at the PM enable the visualization of individual vesicle secretion events and make them an excellent cargo for studying exocytosis in plants. We observed that in cells with reduced myosin or actin activity, many CESA delivery compartments exhibited prolonged pausing at the PM and/or failure to deliver a functional CSC into the PM. In addition, we detected a marked increase in the retention or accumulation of small CESA compartments in the cortical cytoplasm which likely results from failed insertion events. The secretion defects correlated with a reduced delivery rate and decreased density of CSCs at the PM. These new findings reveal a specific role of actin and myosin in vesicle exocytosis, possibly by regulating vesicle tethering or fusion (Fig. 8).

**Figure 8.**
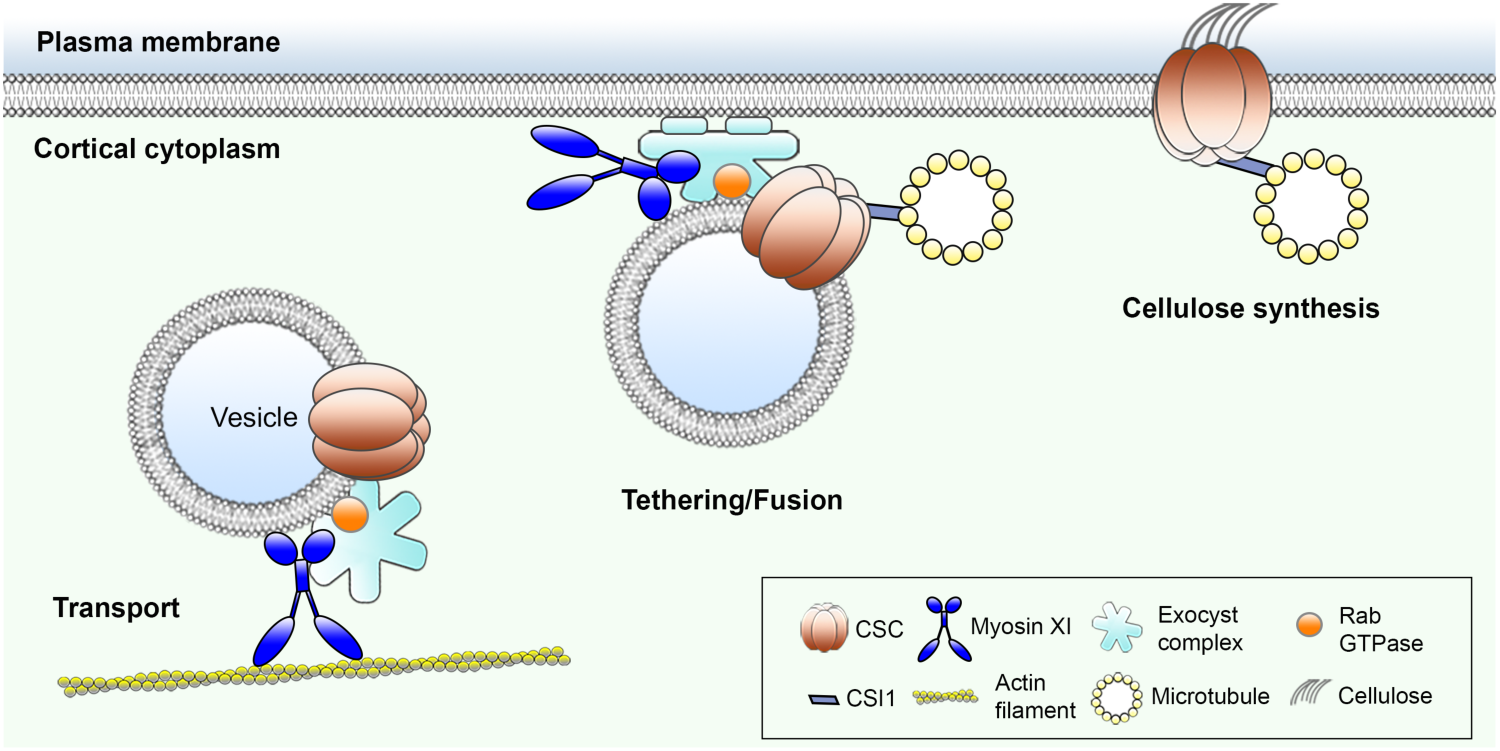
Myosin plays a novel role in vesicle tethering or fusion. This cartoon portrays the known and hypothetical roles of myosin XI in delivery of CSCs to the plasma membrane. Based on evidence herein, as well as recent publications, we predict that myosin not only transports CESA-containing vesicles in the cell cortex, but also participates in vesicle tethering or fusion after vesicles reach the CSC insertion sites that are defined by CSI1 and microtubules. The potential role of myosin in exocytosis could be through interaction with the exocyst complex to stabilize or maintain an intact tethering complex that are required for fusion with the membrane.

In animal and fungal cells, myosin motors have well established roles in secretory vesicle trafficking. They not only transport vesicles to their secretion sites but also participate in the subsequent exocytosis process. Yeast myosin V, Myo2p, has been shown to regulate tethering of secretory vesicles to the PM before fusion and the tethering time is significantly extended in a *myo2p* mutant (Donovan and Bretscher, 2015). In our high-resolution secretion assays, we observed an increased frequency of CESA vesicles that paused longer at the PM in both the *xi3KO* mutant and upon acute myosin inhibitor treatment. It seems likely that myosin XI regulates the tethering of CESA vesicles at the PM, although the detailed mechanism remains to be elucidated. In addition, yeast Myo2p interacts directly with the Sec15 subunit of the exocyst tethering complex and disruption of this interaction blocks secretion and causes significant accumulation of secretory vesicles within the cell (Jin et al., 2011). Considering that the cargo-binding domains of myosins XI are homologous to those of myosin V (Sattarzadeh et al., 2013), and the exocyst shares highly conserved subunit composition and biological functions across eukaryotes (Hála et al., 2008; Synek et al., 2014), plant myosins XI are likely to have conserved function through interaction with exocyst components to regulate vesicle exocytosis. There is limited evidence in plant cells suggesting the interaction of myosin with tethering components; for example, *Arabidopsis* Myosin XIK co-fractionated with the exocyst marker Sec6 in root extracts (Peremyslov et al., 2012). The actin cytoskeleton cooperates with the exocyst subunit EXO84B to execute post-Golgi trafficking of the ATP-binding cassette transporter PENETRATION 3 (PEN3) to the outer periclinal membrane of root epidermal cells (Mao et al., 2016); however, myosin involvement in this polar trafficking has not yet been established.

Our finding of increased number of cortical vesicles in the *xi3KO* mutant or inhibitor-treated cells further supports the hypothesis that myosin XI is involved in vesicle tethering. In plants, the phenotype of accumulation of cortical vesicles is a typical feature found in mutants that are deficient in vesicle tethering. For example, mutations in the exocyst tethering complex, *exo84b,* and the TRAPPII tethering complex, *trs120* and *trs130,* show abnormal accumulation of vesicles in the cortical cytoplasm in *Arabidopsis* (Fendrych et al., 2010; Qi et al., 2011). Although we cannot rule out the possibility that some of the accumulated compartments in myosin-or actin-deficient cells resulted from endocytosis or endocytic vesicles whose intracellular trafficking was interrupted, our studies on internalization of FM4-64 and FLS2 suggest that endocytosis is inhibited in cells with reduced actin and/or myosin activity. In addition, another study shows that following 10-min treatment with mannitol, which increases the number of cortical CESA compartments, the FM4-64 endosomes did not coincide with the CESA compartments, indicating that these vesicles are unlikely to be newly internalized compartments (Gutierrez et al., 2009). Combining the observations of high frequency of vesicle secretion failure and reduced CSC delivery rate in myosin-inhibited cells, we believe that a majority of the accumulated CESA compartments are secretory vesicles, and their accumulation is due to failed membrane tethering and fusion. An overlapping possibility for their accumulation could be that if these are secretory vesicles that failed to fuse to the PM, they might lack the machinery to be transported away from the cortex for recycling or degradation that is associated with newly-formed endocytic vesicles.

A cortical vesicle accumulation phenotype has also been characterized in plant cytokinesis-defective mutants with impaired membrane trafficking during cell plate formation, such as mutations in *KNOLLE* and *KEULE* encoding syntaxins or syntaxin-regulatory proteins that are required for membrane fusion (Lauber et al., 1997; Waizenegger et al., 2000). Mutations of *STOMATAL CYTOKINESIS DEFECTIVE 1 (SCD1)* exhibit secretory vesicle accumulation in cytokinesis-defective cells as well as perturbed cell plate formation. SCD1 and SCD2 have been shown to regulate post-Golgi vesicle exocytosis to the PM through interaction with RabE1 and the exocyst complex (Falbel et al., 2003; Mayers et al., 2017). Interestingly, a recent study also shows concentration of myosin XIK at the forming cell plate in root and shoot meristem cells and the *xi3KO* mutant displays defects in cell division plane alignment (Abu-Abied et al., 2018). Although it remains unclear how myosins XI are involved in cell plate formation and cytokinesis, our findings that myosins XI play a role in exocytic trafficking provides clues that the class XI myosins may share similar functions by mediating secretory membrane trafficking and/or homotypic vesicle fusion during cell plate formation. Further study is required to define and characterize molecular interactions between actin cytoskeletal components and vesicle trafficking machinery proteins, like the exocyst complex or Rab proteins, in the cortical cytoplasm of plant cells. The new role of myosin XI in exocytosis and protein secretion advances our understanding of how myosin activity is correlated with plant cell growth and could help explain the previously described phenotypes in *myosin xi* mutants, such as reduced cell expansion/elongation rate, polar auxin transport and plant size (Peremyslov et al., 2010; Ojangu et al., 2012; Madison et al., 2015; Ueda et al., 2015; Abu-Abied et al., 2018; Ryan and Nebenführ, 2018).

Our colocalization studies reveal that cortical CESA compartments that accumulate in myosin-or actin-deficient cells are distinct from SYP61 or TGN-related compartments, which also accumulate upon myosin inhibition. Some previous studies indicate that the plant trans-Golgi network/early endosomes (TGN/EEs) are involved in both endocytic and secretory trafficking; SYP61 vesicles derived from the TGN are thought to mediate the delivery of various cargos to the PM (Viotti et al., 2010; Drakakaki et al., 2012). However, with respect to CESA trafficking, the role of TGN in secretion of CSCs has been questioned (Crowell et al., 2009). CESA compartments that co-labeled with the TGN/EE marker VHA-a1-mRFP did not associate with CSC insertion events at the PM and are thought to be mainly endocytic compartments (Crowell et al., 2009). Similarly, our results indicate that the CESA-delivery compartments are a distinct population that are not derived from the TGN, which supports the hypothesis that TGN/EEs are not responsible for exocytic trafficking of CSCs in plants.

Since the primary role of the plant actin-myosin transport network is to power cytoplasmic streaming and intracellular organelle motility, it is possible that the cellulose synthesis and trafficking defects detected in myosin-or actin-deficient cells were mainly due to a general perturbation of organelle and vesicle motility. However, our results suggest that this is unlikely the case. We show that in the *xi3KO* mutant that is known to have severely impaired Golgi motility and cytoplasmic streaming rate (Prokhnevsky et al., 2008; Peremyslov et al., 2010), a large motile population of cortical vesicles close to the PM could be detected and they subsequently paused at the PM for CSC delivery, although many of them failed to deliver a CSC particle. This suggests that even though these vesicles rely on the actin cytoskeleton for long-distance transport, their relatively short and local movement to the PM does not necessarily require actin tracks. Similarly, previous reports show that LatB treatment causes gross clustering of Golgi and that CSCs are delivered only to the nearby region (Gutierrez et al., 2009; Sampathkumar et al., 2013), suggesting that the delivery compartments can still travel a certain distance in the cortex but not too far away from Golgi when the cortical actin network is disrupted. In addition, we showed that in the *xi3KO* mutant and upon acute myosin inhibitor treatment, the global Golgi distribution was not greatly affected and no apparent Golgi clustering was detected. Therefore, the CSC trafficking defects in those cells were unlikely caused by a lack of transport of delivery compartments to the PM, but due to inhibition of exocytosis at the PM.

Unexpectedly, we found that myosin perturbation affects the velocity of CSC translocation in the plane of the PM. Because steady movement of CSCs is thought to be powered by the enzymatic activity of CESAs (Paredez et al., 2006; Diotallevi and Mulder, 2007; Wallace and Somerville, 2015), the role of myosin XI is likely to be indirect. One hypothesis is that inhibition of endocytosis in myosin-deficient cells increases the number of slower moving faulty CSCs compared to fully functional CSCs. Similar inhibition of CSC velocities is detected in the *Arabidopsis twd40-2* or *ap2m twd40-2* double mutants; both proteins are components of the clathrin-mediated endocytosis pathway and are involved in CSC internalization (Bashline et al., 2015). Conversely, a delay or incomplete vesicle fusion, like that postulated for exocyst complex mutants (e.g. *sec5*), could cause the accumulation of faulty CSCs held in proximity to the PM (Zhu et al., 2018). It needs to be noted that in contrast to genetic or chemical perturbation of myosin, treatment of cells with LatB had only a modest or no effect on CSC motility in the plane of PM (Supplemental Table S2), as previously reported for actin disruptions (Sampathkumar et al., 2013). This indicates that although the role of myosin as a molecular motor is largely dependent on the actin cytoskeleton, myosin could have a specific role independent from actin. In addition, we observed that myosin-inhibitor treatment, but not LatB, rapidly decreased CESA density at the PM. Similar to a previous publication (Beck et al., 2012), we found that internalization of FLS2 through receptor-mediated endocytosis was markedly reduced with PBP and BDM treatment, but not by LatB. Collectively, these findings suggest that myosin may have a more critical role than actin in exocytic or endocytic trafficking. What exactly is the myosin-centric function in trafficking requires further investigation but could relate to vesicle fusion/scission at the PM.

## METHODS

### Plant Materials and Growth Conditions

An *Arabidopsis* mutant with *myosin xi1, xi2,* and *xik* knocked out (*xi3KO*) and the *xi3KO* mutant expressing vYFP-fABD2 were characterized previously (Peremyslov et al., 2010; Cai et al., 2014). Homozygous YFP-CESA6 *prc1-1* seeds were generated previously (Paredez et al., 2006; Li et al., 2012) and kindly provided by Ying Gu (Pennsylvania State University). The *xi3KO* mutant was crossed to YFP-CESA6 *prc1-1* and F2 homozygous mutants as well as wild-type siblings in the presence of *prc1-1* were recovered. For XIK motility assays, a construct carrying a genomic copy of the *Myosin XIK* gene tagged with YFP previously described in Peremyslov et al. (2012) was transformed into *xi3KO* mutant plants by Agrobacterium-mediated transformation through floral dip (Zhang et al., 2006). For the YFP-CESA6 and SYP61-CFP double-marked line, YFP-CESA6 plants were crossed with plants expressing SYP61-CFP and F1 plants were used for imaging. Transgenic lines of YFP-TUB5 (Shaw et al., 2003), FLS2-GFP (Beck et al., 2012), PIN2-GFP (Xu and Scheres, 2005), BRI1-GFP (Geldner et al., 2007) and SYP61-CFP (Robert et al., 2008) were described previously. Seeds of plants expressing YFP-Mannosidase I were ordered from the Arabidopsis Biological Resource Center (Ohio State University). The YFP-CESA6 mCherry-TUA5 co-expression line was kindly provided by David W. Ehrhardt (Carnegie Institution for Science).

The *Arabidopsis* seeds were surface sterilized and stratified at 4°C for 3 days on half-strength Murashige and Skoog (MS) medium supplemented with 1% sucrose and 0.8% agar. For light growth, plants were grown under long-day lighting conditions (16 h light/8 h dark) at 21°C. For dark growth, plates were exposed to light for 4 h and then placed vertically and kept at 21°C for 3 days in continuous darkness.

### Drug Treatment

For cellulose content and monosaccharide composition analysis, wild-type seedlings were grown on half-strength MS plates containing 100 nM latrunculin B (LatB, Sigma-Aldrich) or 3 mM butanedione monoxime (BDM; Sigma-Aldrich) for 5 d. For short-term live-cell treatments, seedlings were pre-soaked in drug solutions in 24-well plates in the dark. For CSC delivery assays using FRAP and CSC insertion assay, hypocotyls were mounted in inhibitor solutions and imaged immediately. LatB, Pentabromopseudilin (PBP; Adipogen) and MyoVin-1 (EMD Millipore) were dissolved in DMSO to prepare a 5-mM stock solution. BDM was dissolved in water immediately before use. For propidium iodide (PI) staining, seedlings were pre-soaked for 30 s in 10 µg mL^−1^ PI solution dissolved in water. FM4-64 was dissolved in DMSO and used at a concentration of 20 µM.

### Live-Cell Imaging

For image acquisition, epidermal cells at the apical region of 3-d-old dark-grown hypocotyls were used unless otherwise stated. For excitation of YFP-CESA6, XIK-YFP, YFP-TUB5, vYFP-fABD2, secGFP, and SYP61-CFP, spinning-disk confocal microscopy (SDCM) was performed using a Yokogawa scanner unit CSU-X1-A1 mounted on an Olympus IX-83 microscope, equipped with a 100X 1.45–numerical aperture (NA) UPlanSApo oil objective (Olympus) and an Andor iXon Ultra 897BV EMCCD camera (Andor Technology). YFP, GFP and CFP fluorescence was excited with 514-nm, 488-nm, and 445-nm laser lines and emission collected through 542/27-nm, 525/30-nm, and 479/40-nm filters, respectively. For YFP-Mannosidase I imaging, variable-angle epifluorescence microscopy (VAEM) was performed using a TIRF illuminator on an IX-71 microscope (Olympus) as described in Cai et al. (2014). For CSC density and motility imaging, time-lapse images were collected at the plasma membrane (PM) with a 5-s interval for 5 or 61 frames, respectively. For cortical and subcortical YFP-CESA6 and SYP61-CFP imaging, z-series at 0.2 µm step sizes plus time-lapse with 1.6-s intervals for 35 frames were collected. For cortical and subcortical actin analysis, propidium iodide (PI) was used to label the cell wall and as an indicator of the PM plane. Dual-wavelength imaging was performed and z-series at 0.2 µm step sizes were collected. PIN2-GFP and BRI1-GFP were imaged in 6-d old light-grown roots by SDCM as described above with a 60X 1.42 NA UPlanSApo oil objective (Olympus).

For fluorescence recovery after photobleaching (FRAP) experiments, images were collected with a Zeiss Observer Z.1 microscope, equipped with a Yokogawa CSU-X1 and a 100X 1.46 NA PlanApo objective (Zeiss). Photobleaching was performed with a Vector scanner (Intelligent Imaging Innovations) with a 515-nm laser line at 100% power and 3 ms/scan. Time-lapse images were collected at the PM with a 10-s interval for 76 frames, with photobleaching in a small region (44.2 µm^2^) after the 3^rd^ frame, and recovery monitored for a total of 12 min.

### Image Processing and Quantitative Analysis

Image processing and analysis were performed with ImageJ and FIJI (Schindelin et al., 2012). Image drift was corrected with StackReg plugin. Golgi motility was analyzed with TrackMate plugin and a Laplacian of Gaussian (LoG) algorithm as particle detection filter. Trajectories detected by TrackMate were selected for analysis only if more than 5 spots were on the trajectory. The parameter “Mean Speed” was then pooled, and plotted as the average Golgi motility rate. XIK-YFP motility was measured by Multi Kymograph plugin. For PM-localized CSC density analysis, the first frames of time-lapse images were used and no intensity adjustment was made. The number of CSC was then counted by TrackMate plugin with Difference of Gaussians (DoG) algorithm. CSC density was calculated as the number of particles detected divided by the area of the analyzed region. For PM-localized CSC motility assays, average intensity projections were generated with the time-lapse series to identify the trajectories of the CSC particles over time. Kymographs were generated by following the trajectories of CSC particles and CSC motility rate was calculated as the reciprocal of the slope of individual CSC particles in kymographs. For FRAP assays, a smaller area (28.3 µm^2^) within the bleached region was used for analysis to exclude the lateral movement of CSCs into the bleached region (Sampathkumar et al., 2013). The newly delivered CSCs during the initial 5 min of recovery were manually counted and the criteria used to define a CSC delivery event were as described previously (Gutierrez et al., 2009; Li et al., 2016). Only the newly inserted particles that exhibited a static phase followed by steady linear migration at the PM were counted as new delivery events. The CSC delivery rate was calculated as the number of delivery events divided by the measured area and time. For cortical and subcortical vesicle assays, z-series taken at 0.2 µm and 0.4 µm below the PM were used for cortical cytoplasm analysis, and focal planes from 0.6 µm to 1.0 µm as the subcortical cytoplasm. Image background was subtracted using Subtract Background tool in FIJI with rolling ball radius set at 30 pixels. The number of CESA and SYP61 vesicles was counted manually based on their size (smaller than a Golgi or TGN compartment) and dynamic behavior in the time-lapse series (Gutierrez et al., 2009). To determine overlap between CESA and SYP61 vesicles, only signals from both fluorescence channels that moved in the same trajectory over time were scored as colocalization. Vesicle density was calculated as the number of particles detected divided by the area of the analyzed region. The velocities of CESA compartments were measured by semi-automatic tracking with MTrackJ (Meijering et al., 2012) or Multi Kymograph plugin. The mean velocity of each individual trajectories was used for data analysis. For analysis of individual CSC insertion events, only particles that showed *de novo* appearance at the PM plane followed by a pausing phase for more than 5 frames (> 15 s) were considered as new insertion events. The pausing phase was determined by analysis of kymographs. Only a straight line in the kymograph was considered as a pausing event. The exact duration of particle pausing time was determined by fitting a straight line along the moving trajectory and another line along the pausing phase in the kymograph. The intersection of the two lines was defined as the end of the pausing phase. For analysis of actin filament array organization, actin filament density and the extent of bundling (skewness) were determined as described previously (Higaki et al., 2010; Henty et al., 2011).

### FM4-64 Internalization Assay

Four-day-old light-grown seedlings of wild type and the *xi3KO* mutant were pre-soaked in inhibitor solutions for 30 min followed by incubation in 20 µM FM4-64 plus the inhibitor solutions for 6 min in the dark. Images were collected at the hypocotyl epidermis with SDCM as described above with a 60X 1.42 NA UPlanSApo oil objective (Olympus). The number of internalized FM4-64 endosomes in individual cells was counted and the density was calculated as the number of endosomes per 100 µm^2^.

### FLS2-GFP Internalization Assay

Seven-day-old transgenic seedlings expressing FLS2-GFP were pre-soaked in dH_2_O overnight in continuous light to reduce wounding response as described previously (Smith et al., 2014). Seedlings were then soaked in inhibitor solutions for 30 min followed by treatment with 1 µM flg22 (NeoBioSci) plus the inhibitors for another 45 min to induce the endocytosis of FLS2. Cotyledon epidermal cells at the adaxial side were imaged by SDCM as described above with a 60X 1.42 NA UPlanSApo oil objective (Olympus). Z-series were collected at 0.5-µm step size for 21 frames. Maximum intensity projections were generated from the z-series and quantitative analysis of internalized FLS2-GFP endosomes was done using Advanced Weka Segmentation plugin in FIJI as described in Smith et al. (2014).

### Cytoplasmic Streaming Assay of *Chara corallina*

Young internodal cells of *C. corallina* less than 30 mm long were cut into ∼7 mm pieces and soaked in the original culture water containing drugs in 24-well plates at room temperature. Light microscopy was performed using a Nikon E600 microscope with a 20X 0.5 NA Plan Fluor objective. Time-lapse images at 0.22-s intervals were collected with a CCD camera (ORCA-ER C4742-95; Hamamatsu Photonics) and Metamorph software (version 4.6r9). The velocity of individual particles was analyzed by manual tracking with MTrackJ plugin in FIJI. The mean velocities measured by the plugin were pooled and used for statistical analysis.

### Cell Wall Determination

Five-d-old dark-grown hypocotyls were lyophilized and ground in SDS buffer (1% SDS, 50 mM Tris-HCl, pH 7.2). The homogenate was washed sequentially with SDS buffer and 50% ethanol at 60°C, and washed with acetone at room temperature. The acetone was discarded and samples were vacuum dried overnight. The dry residue was defined as cell wall material (CWM).

The cellulose content was determined as modified from Updegraff (1969). Briefly, about 2-3 mg of CWM was hydrolyzed with acetic-nitric reagent (acetic acid:nitric acid:water = 8:1:2) at 100°C for 60 min; the insoluble material was collected after centrifugation at 2500 × *g* for 5 min, and washed with water. Alternatively, about 2.5 mg of CWM was hydrolyzed in 2 M trifluoroacetic acid (TFA) containing 500 nmole mL^−1^ of *myo*-inositol (internal standard) at 120°C for 90 min. The insoluble material was collected by centrifugation as above, and the soluble fraction saved for non-cellulosic monosaccharide analysis. The amounts of sugar in the insoluble fractions from both the acetic-nitric and TFA hydrolysis were measured using phenol-sulfuric colorimetric assay compared to cellulose standards (Dubois et al., 1956).

Non-cellulosic monosaccharide composition in the supernatant fractions from TFA hydrolysis were determined by separation and quantitation of alditol acetates by GLC-MS. Briefly, the TFA was evaporated in a stream of nitrogen gas in the presence of *tert*-butyl alcohol to minimize destruction in concentrated acid, and monosaccharides with reduced with NaBH_4_ and acetylated with acetic anhydride (Gibeaut and Carpita, 1991). The derivatives were separated into seven components representing the major sugars in plant cell walls by gas-liquid chromatography on an SP-2330 (Supelco, Bellefonte, PA) using a 0.25-mm × 30-m column with temperatures from 80°C to 170°C increased 25°C min^−1^ followed by an increase of 5°C min^−1^ up to 240°C and a helium flow of 1 mL min^−1^. Electron impact mass spectrometry was carried out on a Hewlett-Packard MSD at 70 eV with a source temperature of 250°C. Area under the curve calculations for each sugar derivative were scaled to µg per mg of sample tissue using the *myo*-inositol internal standard (Carpita and Shea, 1989).

### Accession Numbers

Sequence data from this article can be found in the Arabidopsis Genome Initiative under the following accession numbers: *Myosin XIK*, At5g20490; *Myosin XI1*, At1g17580; and *Myosin XI2*, At5g43900.

## Supporting information

## Supplemental Material

The following materials are available in the online version of this article.

**Supplemental Table 1.** Cell wall monosaccharide composition of wild-type, *xi3KO* mutant, and LatB-treated seedlings.

**Supplemental Table 2.** Summary of parameters affected by genetic or chemical disruption of plant myosin XI.

**Supplemental Figure 1.** Cellulose content is reduced in BDM-treated seedlings.

**Supplemental Figure 2.** Distribution of individual Golgi velocities following treatment with myosin inhibitors.

**Supplemental Figure 3.** PBP treatment reduces Golgi and myosin XIK motility in a dose-and time-dependent manner and is reversible.

**Supplemental Figure 4.** PBP inhibits cytoplasmic streaming in *Chara corallina* internodal cells.

**Supplemental Figure 5.** Disruption of myosin does not cause clustering of CESA-containing Golgi.

**Supplemental Figure 6.** Selected regions of interest from control and inhibitor-treated cells have the same density of PM-localized CSCs at the beginning of FRAP experiments.

**Supplemental Figure 7.** Disruption of cortical microtubules does not alter the rate of CSC delivery to the PM.

**Supplemental Figure 8.** Inhibition of myosin activity reduces delivery of PIN2-GFP and BRI-GFP to the PM and leads to retention of intracellular secGFP.

**Supplemental Figure 9.** Velocity of CESA compartments is reduced in both cortical and subcortical cytoplasm of myosin-deficient cells.

**Supplemental Figure 10.** Internalization of FM4-64 and FLS2 is inhibited in myosin-deficient cells.

**Supplemental Figure 11.** Actin architecture in the cortical and subcortical cytoplasm of *xi3KO* mutant and myosin inhibitor-treated cells is altered.

**Supplemental Figure 12.** Inhibition of myosin results in accumulation of SYP61 vesicles in the cortical cytoplasm.

**Supplemental Figure 13.** Inhibition of myosin does not affect the microtubule-dependent positioning of CSC insertion sites.

**Supplemental Movie 1.** Myosin inhibitor treatments reduce Golgi motility.

**Supplemental Movie 2.** Myosin inhibitor treatments reduce myosin XIK motility.

**Supplemental Movie 3.** Small CESA compartments accumulate in the cortical cytoplasm of myosin-and actin-deficient cells.

**Supplemental Movie 4.** A successful CSC insertion event at the PM.

**Supplemental Movie 5.** A CSC particle is inserted next to a cortical microtubule and then migrates along the microtubule.

**Supplemental Movie 6.** A failed CSC insertion event that had a short pausing time at the PM.

**Supplemental Movie 7.** A failed CSC insertion event that had a long pausing time at the PM.

**Supplemental Movie Legends.** Legends for Supplemental Movies 1 to 7.

## ACKNOWLEDGEMENTS

We thank Nick Carpita and Anna Olek (Purdue) for assistance and access to equipment for cellulose determinations. Cellulose and monosaccharide analyses were supported by the Center for the Direct Catalytic Conversion of Biomass to Biofuels, an Energy Frontiers Research Center of the U.S. Department of Energy, Office of Science, Basic Energy Sciences (grant no. DE–SC0000997). We thank Valerian Dolja (Oregon State University) for providing the *myosin xi* triple knockout, Ying Gu (Penn State University) for sharing the CESA6-YFP complementation line, and Chunhua Zhang (Purdue) for helpful discussions. The authors are grateful to Hongbing Luo (Purdue) for excellent care and maintenance of plant materials.

