## Supplementary material for "*Arabidopsis* Myosins XI Are Involved in Exocytosis of Cellulose Synthase Complexes"

**Supplemental Data**

**Supplemental Table 1.** Cell wall monosaccharide composition of wild-type, *xi3KO* mutant, and LatB-treated seedlings.

| | Rha | Fuc | Ara | Xyl | Man | Gal | Gluc |
|---|---|---|---|---|---|---|---|
| WT | 24.7 ± 0.9[‡] | 4.8 ± 0.3 | 34.8 ± 2.2 | 49.7 ± 3.1 | 8.6 ± 0.8 | 63.7 ± 2.9 | 43.6 ± 1.5 |
| LatB | 28.9 ± 2.0 | 4.7 ± 0.3 | 44.9 ± 2.9* | 44.9 ± 2.7 | 8.1 ± 0.4 | 67.6 ± 4.0 | 37.5 ± 2.2 |
| *xi3KO* | 24.6 ± 1.4 | 4.4 ± 0.2 | 35.9 ± 2.1 | 42.0 ± 2.3 | 8.4 ± 0.6 | 55.9 ± 3.5 | 42.5 ± 2.7 |

[‡]Monosaccharides in µg $mg^{-1}$ of cell wall material; Rha = Rhamnose; Fuc = Fucose; Ara = Arabinose; Xyl = Xylose; Man = Mannose; Gal = Galactose; Gluc = Glucose; Values are means ± SE of 4 biological replicates; Student's *t* test, $*P < 0.05$

**Supplemental Table 2.** Summary of parameters affected by genetic or chemical disruption of plant myosin XI

| Genotype /Treatment | Cellulose content [A] | Golgi motility [B] | XIK motility [B] | CSC density at the PM [C] | CSC motility at PM [C] | CSC delivery rate [D] | Cortical CESA comp. density [E] | Subcortical CESA comp. density [E] | Cortical SYP61 comp. density [F] | Frequency of failed insertion [G] | CSC pausing time at the PM [G] | PIN2-GFP at the PM [H] | BRI1-GFP at the PM [H] | Intracellular secGFP [H] | FM4-64 internalization [I] | FLS2-GFP internalization [I] | Cortical actin density [J] | Cortical actin bundling [J] |
|---|---|---|---|---|---|---|---|---|---|---|---|---|---|---|---|---|---|---|
| ***xi3KO*** | ↓ | – | – | ↓ | ↓ | ↓ | ↑ | ND | – | ↑ | ↑ | – | – | – | ↓ | – | ↓ | ↑ |
| **BDM** | ↓ | ↓ | ↓ | ↓ | ↓ | ↓ | ↑ | ND | ↑ | ↑ | ↑ | ↓ | ↓ | ↑ | ↓ | ↓ | ↑ | ↓ |
| **PBP** | – | ↓ | ↓ | ↓ | ↓ | ↓ | ↑ | ↑ | ↑ | ↑ | ↑ | ↓ | ↓ | ↑ | ↓ | ↓ | ↓ | ↑ |
| **MyoVin-1** | – | ND | ND | – | – | – | – | – | – | – | – | – | – | – | – | – | – | – |
| **LatB** | ↓ | – | – | ND | ND | ↓ | ↑ | ND | ↑ | ↑ | ↑ | ↓ | ↓ | ↑ | ↓ | ND | ↓ | ↑ |

↓ = decrease in value; ↑ = increase in value; ND = no difference; comp. = compartment

A: see Figure 1; B: see Figure 2; C: see Figure 3; D: see Figure 4; E: see Figure 5; F: see Supplemental Figure 12; G: see Figure 7; H: see Supplemental Figure 8; I: see Supplemental Figure 10; J: see Supplemental Figure 11.

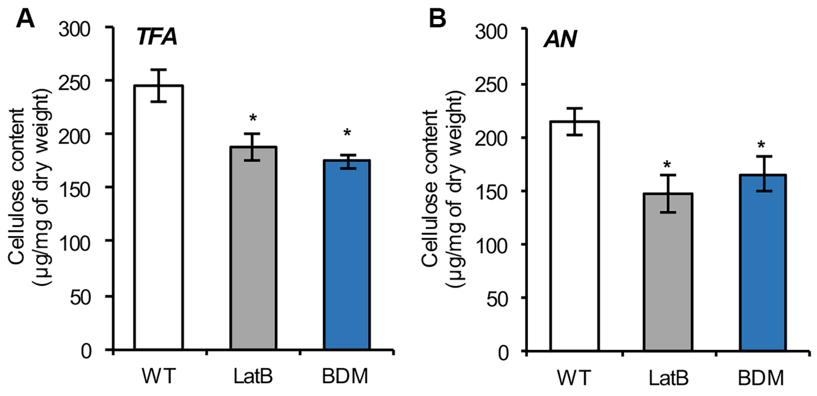


**Supplemental Figure 1.** Cellulose content is reduced in BDM-treated seedlings. Ethanol-insoluble cell wall material (CWM) was prepared from 5-d-old etiolated hypocotyls of wild-type (WT) seedlings, as well as WT growing on plates containing 100 nM latrunculin B (LatB) or 3 mM 2,3-butanedione monoxime (BDM).

A and B, The non-cellulosic component of CWM was hydrolyzed with 2 M trifluoroacetic acid (TFA; A) for total cellulose determination or acetic nitric reagent (AN; B) for crystalline cellulose determination. Cellulose content was significantly reduced in LatB- and BDM-treated seedlings compared to WT. Values given are means ± SE (n = 4; Student's *t* test, $*P < 0.05$).

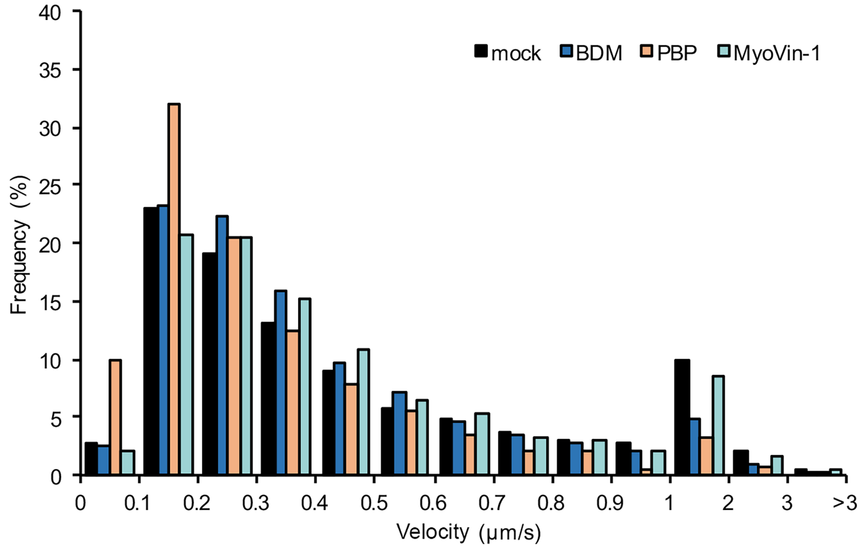


**Supplemental Figure 2.** Distribution of individual Golgi velocities following treatment with myosin inhibitors.

Compared with mock treatment, PBP treatment resulted in a higher proportion of Golgi that move at <0.2 µm $s^{-1}$ and fewer Golgi that moved at >0.6 µm $s^{-1}$. In contrast, BDM treatment mainly reduced the proportion of Golgi with a velocity >1 µm $s^{-1}$. Data are from the same experiment shown in Fig. 2C (n > 3000 trajectories from 10 hypocotyls per treatment).

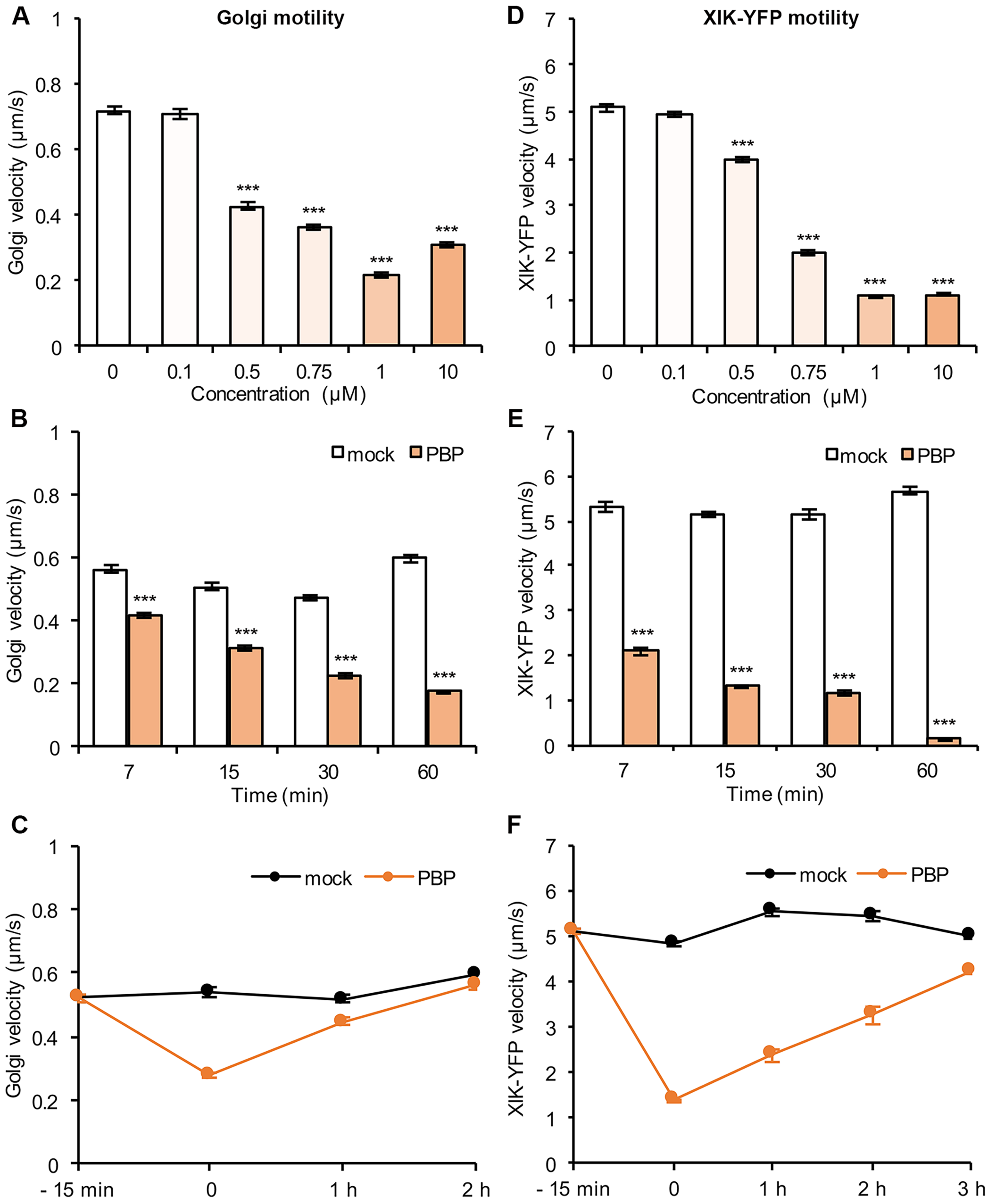


**Supplemental Figure 3.** PBP treatment reduces Golgi and myosin XIK motility in a dose- and time-dependent manner and is reversible.

A and D, Quantification of Golgi and myosin velocity in apical hypocotyl epidermal cells expressing YFP-Mannosidase I and Myosin XIK-YFP, respectively. Seedlings were treated with mock (0.2% DMSO), 0.1, 0.5, 0.75, 1 and 10 μM PBP for 15 min prior to imaging. The velocity of Golgi and XIK-YFP movement was significantly decreased in a dose-dependent fashion. B and E, Hypocotyls were treated with mock (0.2% DMSO) or 10 μM PBP for 7, 15, 30 and 60 min before imaging. The decrease of Golgi and XIK-YFP velocity with PBP treatment was time-dependent. C and F, Hypocotyls were imaged before treatment, after treatment with mock (0.2% DMSO) or 1 μM PBP for 15 min, and after wash out of PBP for 1, 2 and 3 h. The motility of Golgi and XIK-YFP gradually recovered reaching mock levels after wash out of PBP for 2–3 h. Values given are means ± SE. For Golgi velocity measurements, n > 2000 trajectories from 8–12 hypocotyls per treatment; for XIK-YFP velocity measurements, n > 300 tracks from 8–12 hypocotyls per treatment; Student's t test, ***P < 0.001.

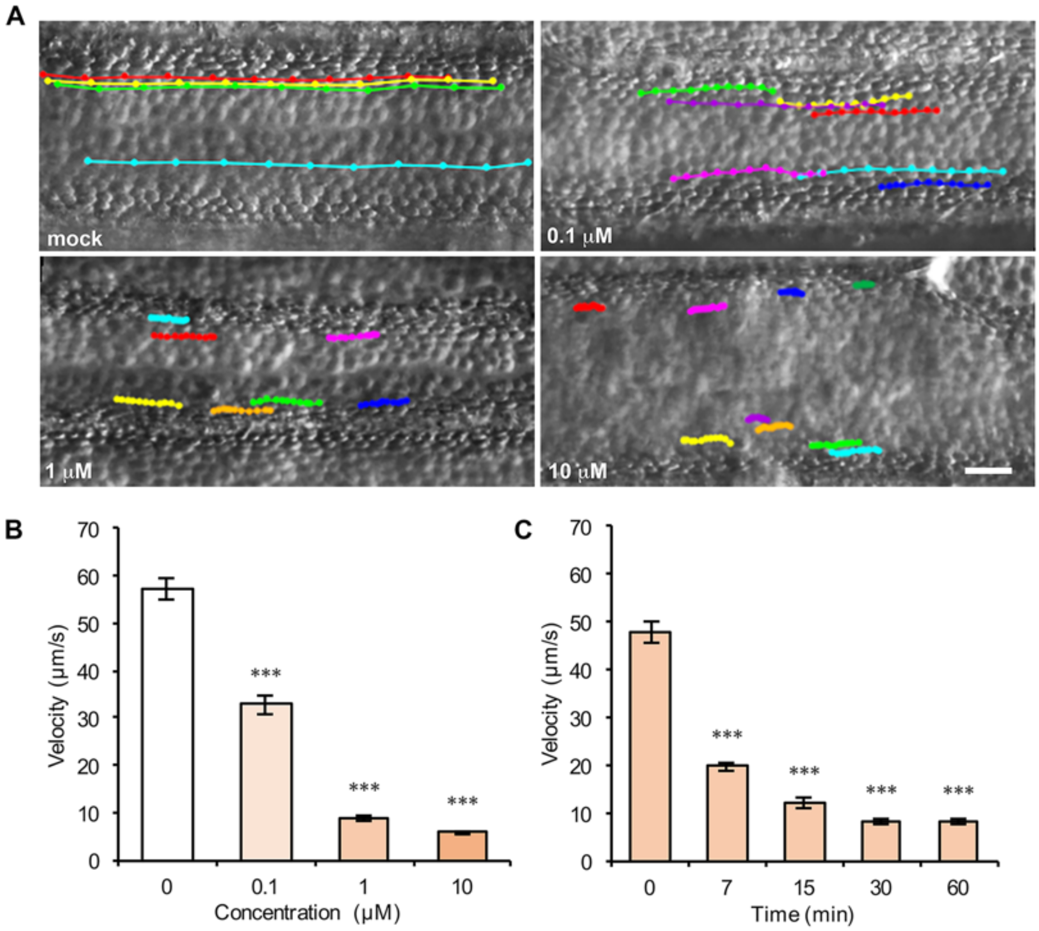


**Supplemental Figure 4.** PBP inhibits cytoplasmic streaming in *Chara corallina* internodal cells.

A, Representative light micrographs and trajectories of moving particles in the cytoplasm of *C. corallina* internodal cells treated with mock (0.2% DMSO), 0.1, 1 and 10 μM PBP for 30 min prior to imaging. Trajectories were generated using time-lapse images taken at 0.22 s-intervals for a total of 2.2 s. The trajectories were much shorter in PBP-treated cells compared with those in mock-treated cells. Bar = 10 μm. B, Analysis of average particle velocity showed that cytoplasmic streaming was significantly inhibited by PBP in a dose-dependent fashion. C, Cells were treated with 1 μM PBP for 0, 7, 15, 30 and 60 min before imaging. The inhibition of cytoplasmic streaming velocity with PBP treatment was time-dependent. Values given are means ± SE (n > 50 particles from 10 cells per treatment, Student's *t* test, *** P < 0.001).

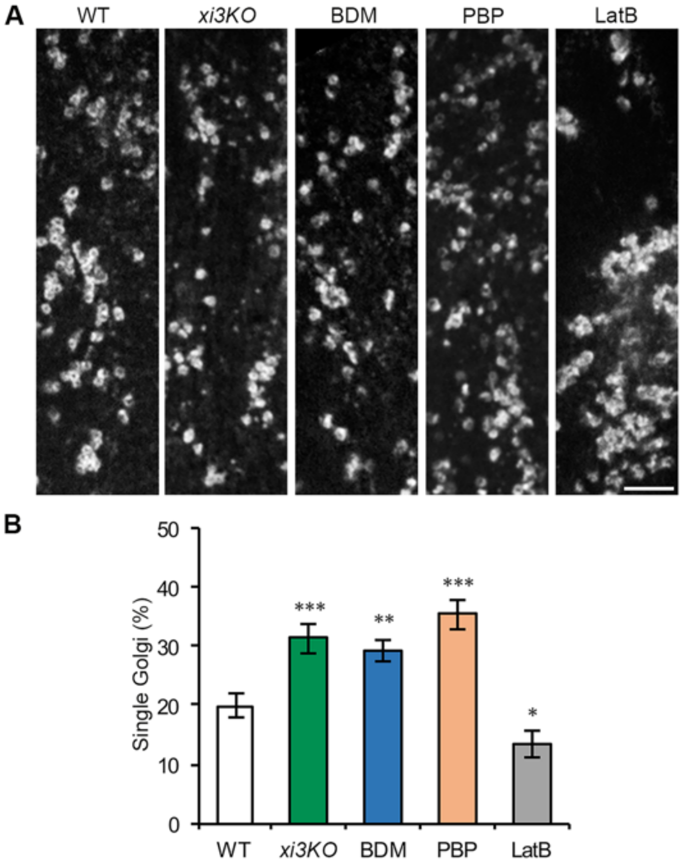


**Supplemental Figure 5.** Disruption of myosin does not cause clustering of CESA-containing Golgi.

A, Representative single optical sections from the cortical cytoplasm of dark-grown hypocotyl epidermal cells expressing YFP-CESA6. Seedlings of *xi3KO* or its WT sibling were treated with mock (0.2% DMSO), 30 mM BDM, 10 μM PBP or 10 μM LatB for 15 min prior to imaging. Compared with WT, a clustering of Golgi was observed in LatB-treated cells, whereas in *xi3KO*, BDM- and PBP-treated cells, Golgi were more evenly distributed. Bar = 5 μm. B, Quantification of the percentage of single unclustered Golgi. The percentage of single Golgi increased significantly in *xi3KO*, BDM- and PBP-treated cells, but was reduced in LatB-treated cells compared with WT. Values given are means ± SE (n ≥ 23 cells from 10 hypocotyls for each genotype or treatment, Student's *t* test, ***P < 0.001, **P < 0.01, *P < 0.05).

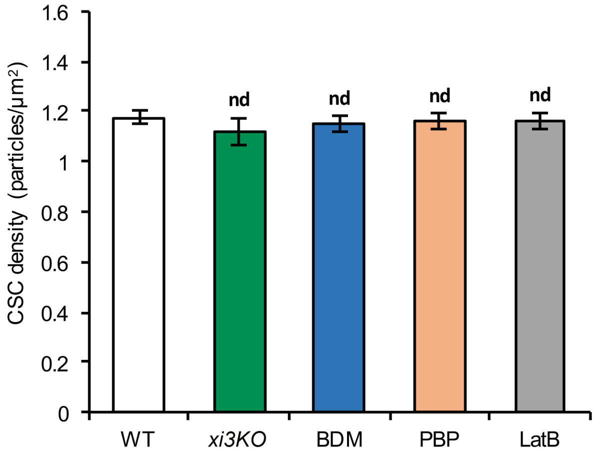


**Supplemental Figure 6.** Selected regions of interest from control and inhibitor-treated cells have the same density of PM-localized CSCs at the beginning of FRAP experiments.

The density of CSCs in regions of interest selected for photobleaching was similar in the *xi3KO* mutant, BDM-, PBP- and LatB-treated cells compared with WT. Data are from the same experiment shown in Fig. 4B. Values given are means ± SE (n = 9–11 cells from 9–11 hypocotyls per genotype or treatment; Student's *t* test, nd: $P > 0.05$).

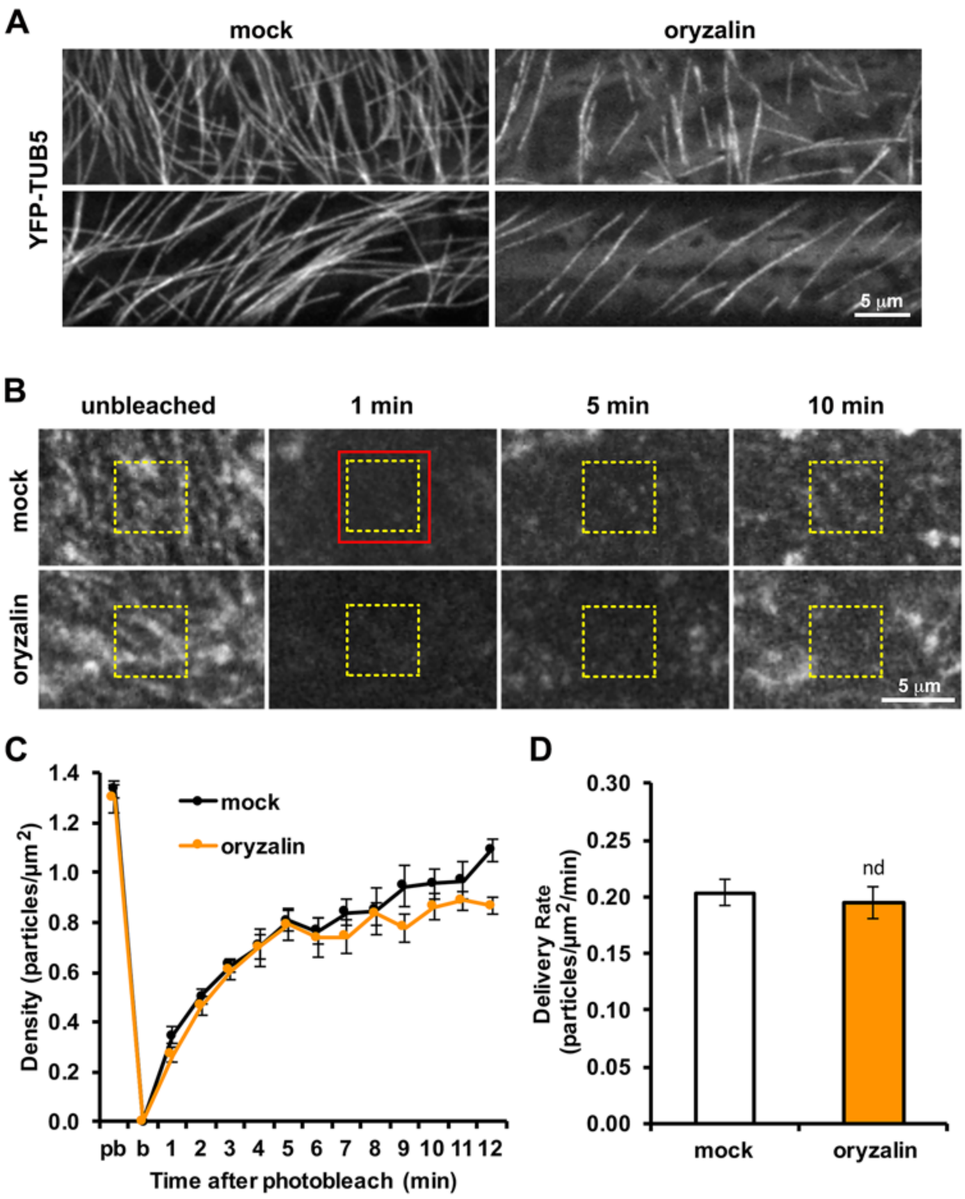


**Supplemental Figure 7.** Disruption of cortical microtubules does not alter the rate of CSC delivery to the PM.

A, Representative images show the organization of cortical microtubules in epidermal cells of 3-d-old dark-grown hypocotyls expressing YFP-TUB5 following treatment with mock (0.1% DMSO) or 20 µM oryzalin for 2 h. B, Representative single-frame images of PM-localized CSC particle recovery after photobleaching. Seedlings expressing YFP-CESA6 were pre-treated with mock (0.1% DMSO) or 20 µM oryzalin for 2 h. A region of interest at the PM (red box) was photobleached and a subarea within the region (yellow dashed box) was used for analysis of particle recovery. C, The recovery of CSCs at the PM after photobleaching over a 12-min period. D, The rate of delivery of CSCs to the PM was calculated as the slope of linear regression from plots of CSC density during the initial 3 min of recovery. The delivery rate was not significantly different between mock- and oryzalin-treated cells. Values given are means ± SE (n = 10 cells per treatment, Student's *t* test, nd: P > 0.05).

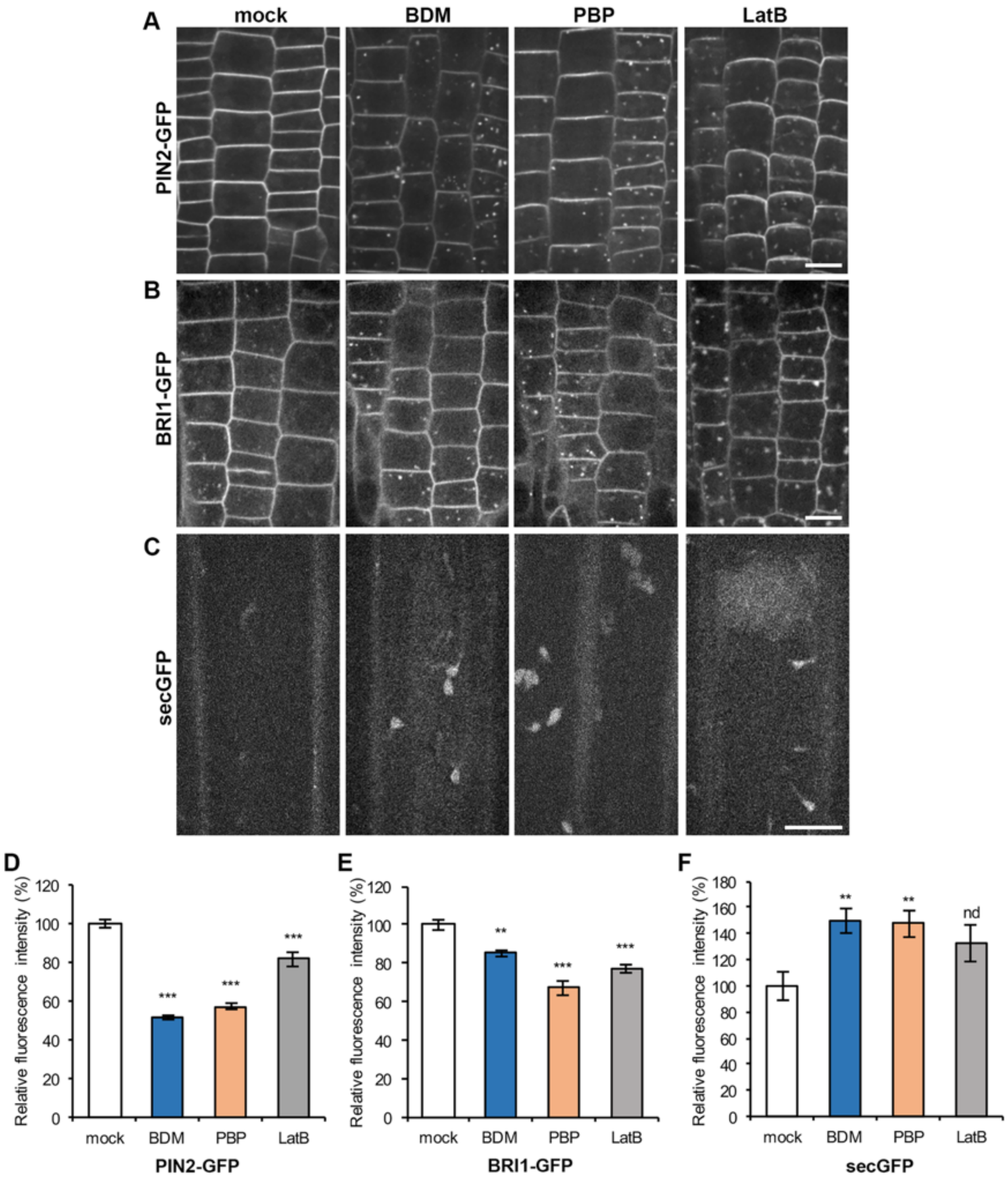


**Supplemental Figure 8.** Inhibition of myosin activity reduces delivery of PIN2-GFP and BRI-GFP to the PM and leads to retention of intracellular secGFP.

A and B, Representative images of 6-d old root epidermal cells expressing PIN2-GFP (A) or BRI1-GFP (B). Seedlings were treated with mock (0.2% DMSO), 30 mM BDM, 10 µM PBP or 10 µM LatB for 2 h. Bars = 10 µm. C, Representative images show cortical cytoplasm of 3-d old etiolated hypocotyls cells expressing secGFP. Seedlings were treated in inhibitor solutions for 30 min. Bar = 10 µm. D and E, Quantification shows that the fluorescence intensity of PIN2-GFP (D) and BRI1-GFP (E) at the PM was greatly reduced after BDM-, PBP- and LatB-treatment. Values given are means ± SE (n ≥ 29 cells from 5 seedlings per treatment; Student's *t* test, **P < 0.01, ***P < 0.001). F, A significantly higher amount of intracellular secGFP fluorescence was retained in BDM- and PBP-treated cells compared to mock treatment. Values given are means ± SE (n ≥ 27 images from 6 seedlings per treatment; Student's *t* test, **P < 0.01, nd: P > 0.05).

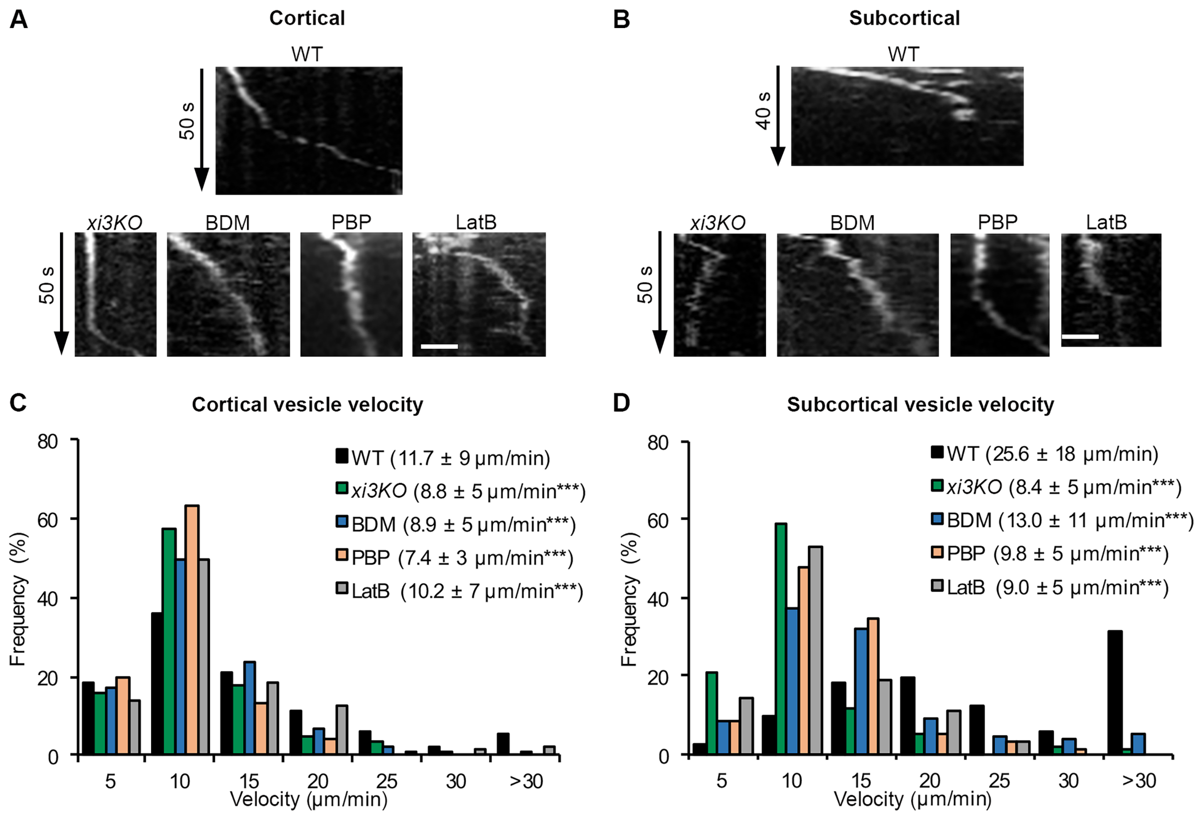


**Supplemental Figure 9.** Velocity of CESA compartments is reduced in both cortical and subcortical cytoplasm of myosin-deficient cells.

A and B, Representative kymographs show the movement of CESA compartments at the cortical (A) and subcortical (B) cytoplasm of epidermal cells from *xi3KO* or WT siblings expressing YFP-CESA6 treated with mock (0.2% DMSO), 30 mM BDM, 10 µM PBP or 10 µM LatB for 15 min. CESA compartments moved shorter distances in both the cortical and subcortical cytoplasm during a 50 s interval in *xi3KO*, BDM-, PBP- or LatB-treated cells compared with WT. Bars = 2 µm. C and D, Histograms of the average velocities of CESA compartments in the cell cortex (C) and subcortex (D). The mean velocities in *xi3KO* or BDM-treated cells were significantly reduced compared with WT. Values given are means ± SD (For cortical vesicles, n = 168, 185, 145, 123 and 173 particles for WT, *xi3KO*, BDM, PBP and LatB, respectively; for subcortical vesicles, n = 82, 92, 110, 61 and 64 particles for WT, *xi3KO*, BDM, PBP and LatB, respectively; Student's *t* test, ***P < 0.001).

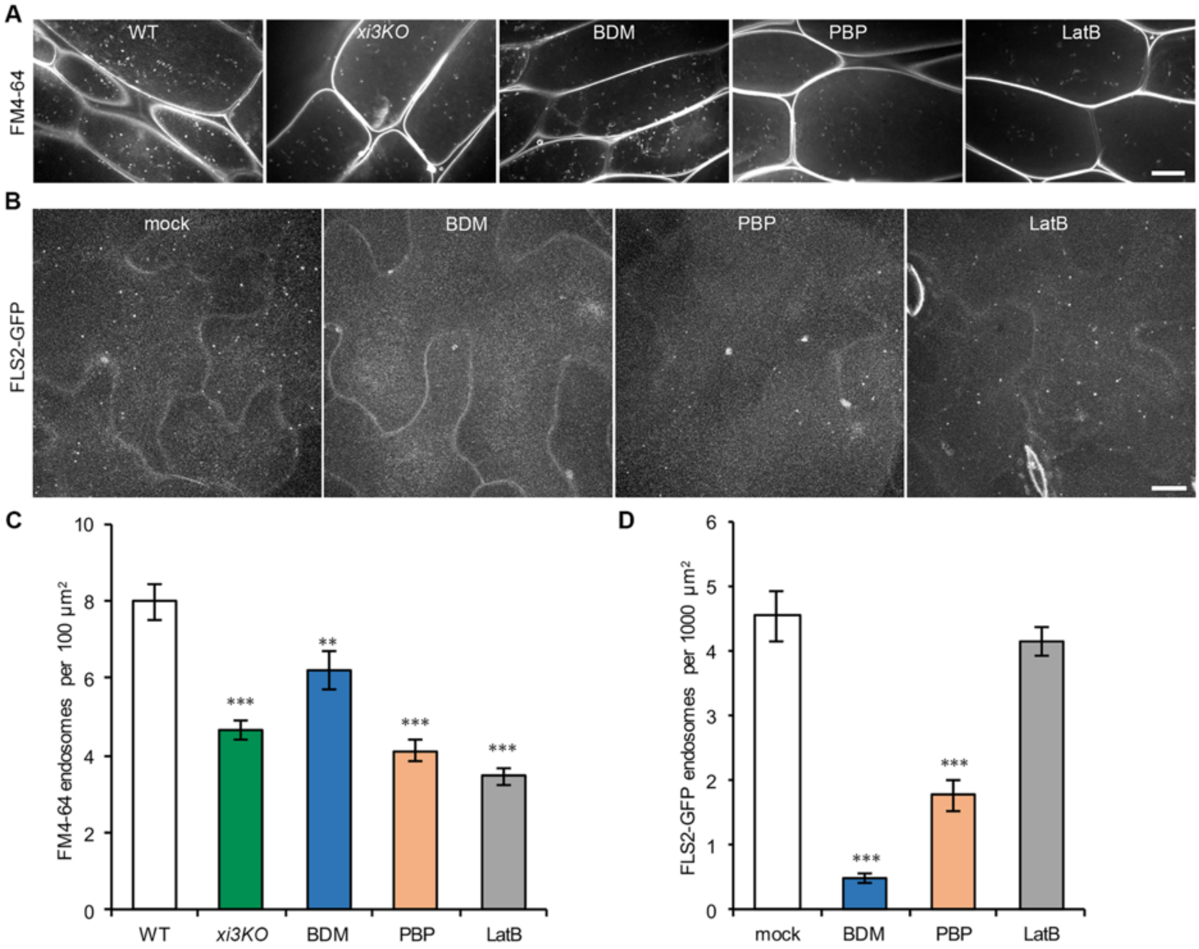


**Supplemental Figure 10.** Internalization of FM4-64 and FLS2 is inhibited in myosin-deficient cells.

A, Representative single optical sections show internalization of FM4-64 endosomes in hypocotyl epidermis. Four-d-old light grown seedlings of *xi3KO* or WT were treated with mock (0.2% DMSO), 30 mM BDM, 10 µM PBP or 10 µM LatB for 30 min followed by addition of 20 µM FM4-64 for 6 min. Bar = 10 µm. B, Representative maximum intensity projections of 7-d-old *Arabidopsi*s cotyledon epidermal cells expressing FLS2-GFP after treatment with flg22 for 45 min. Seedlings were pre-treated with mock (0.2% DMSO), 30 mM BDM, 10 µM PBP or 10 µM LatB for 30 min prior to flg22 treatment. Bar = 10 µm. C, Quantitative analysis shows that the internalization of FM4-64 was significantly inhibited in *xi3KO*, PBP-, BDM- and LatB-treated cells. Values given are means ± SE (n ≥ 60 cells from 8 hypocotyls per treatment; Student's *t* test, ***P < 0.001). D, Quantification shows that BDM and PBP treatment significantly reduced the number of FLS2 endosomes. Values given are means ± SE (n ≥ 55 images from 6-12 seedlings per treatment; Student's *t* test, ***P < 0.001).

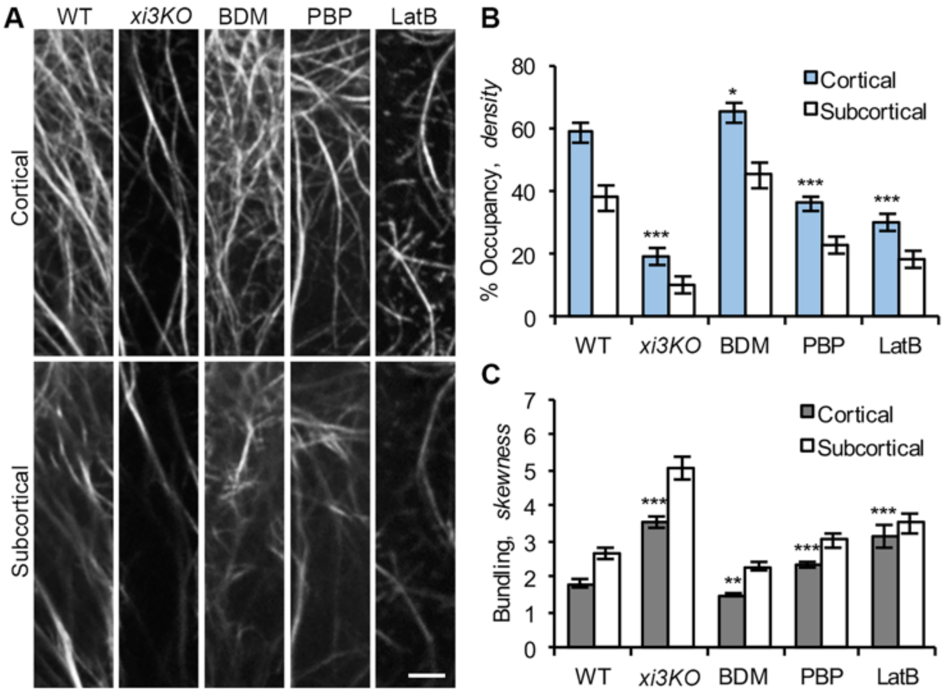


**Supplemental Figure 11.** Actin architecture in the cortical and subcortical cytoplasm of *xi3KO* mutant and myosin inhibitor-treated cells is altered.

A, Representative single optical sections from epidermal cells of 3-d-old dark-grown hypocotyls expressing vYFP-fABD2. Seedlings of *xi3KO* or WT siblings were treated with mock (0.2% DMSO), 30 mM BDM, 10 μM PBP or 10 μM LatB for 15 min prior to imaging. Images were collected at 0.2 μm or 0.8 μm below the PM, which represent the cortical and subcortical cytoplasm, respectively. Bar = 5 μm. B and C, Quantitative analyses of the density and the extent of bundling of actin arrays in the cortical and subcortical focal planes. The overall patterns of cortical and subcortical actin architecture in *xi3KO*, PBP- and BDM-treated cells were similar to those in WT. However, *xi3KO* and PBP treatment had reduced density and increased bundling compared to WT, whereas BDM-treated cells had increased density and reduced bundling. Values given are means ± SE (n > 50 cells from 15 hypocotyls for each genotype or treatment, Student's *t* test, *P < 0.05, **P < 0.01, ***P < 0.001).

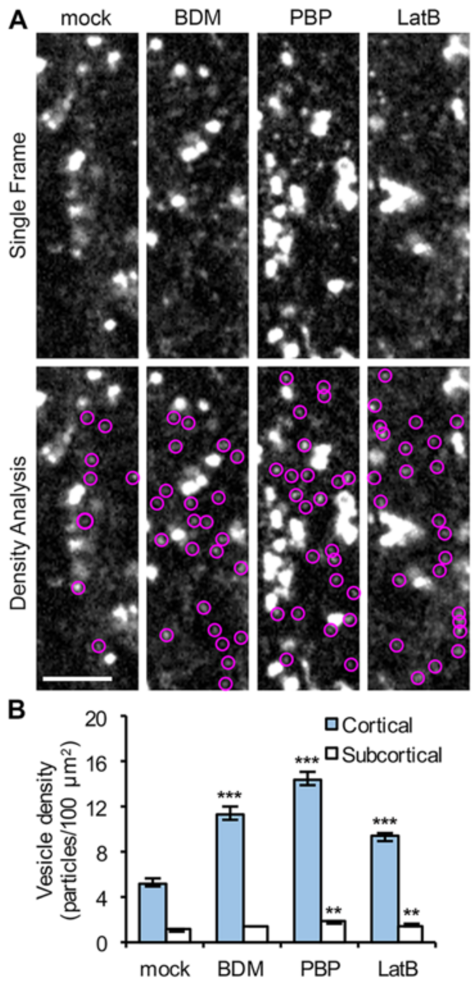


**Supplemental Figure 12.** Inhibition of myosin results in accumulation of SYP61 vesicles in the cortical cytoplasm.

A, Representative single frames taken at the cortical focal plane (0.4 μm below the PM) in etiolated hypocotyl epidermal cells expressing SYP61-CFP. Seedlings were treated in inhibitor solutions for 15 min prior to imaging. The SYP61 vesicles are highlighted with magenta circles. Bar = 5 μm. B, Quantitative analysis of vesicle density shows that the number of SYP61 vesicles was increased significantly after BDM, PBP and LatB treatment and the vesicles were preferentially accumulated in the cortical cytoplasm rather than the subcortical cytoplasm. Values given are means ± SE (n ≥ 30 cells from 10 seedlings for each genotype or treatment; a total number of 493, 971, 1184 and 768 compartments were measured in total areas of 7754, 7545, 7316 and 7088 $\mu m^2$ in mock-, BDM-, PBP- and LatB-treated cells, respectively. Student's *t* test, **P < 0.01, ***P < 0.001).

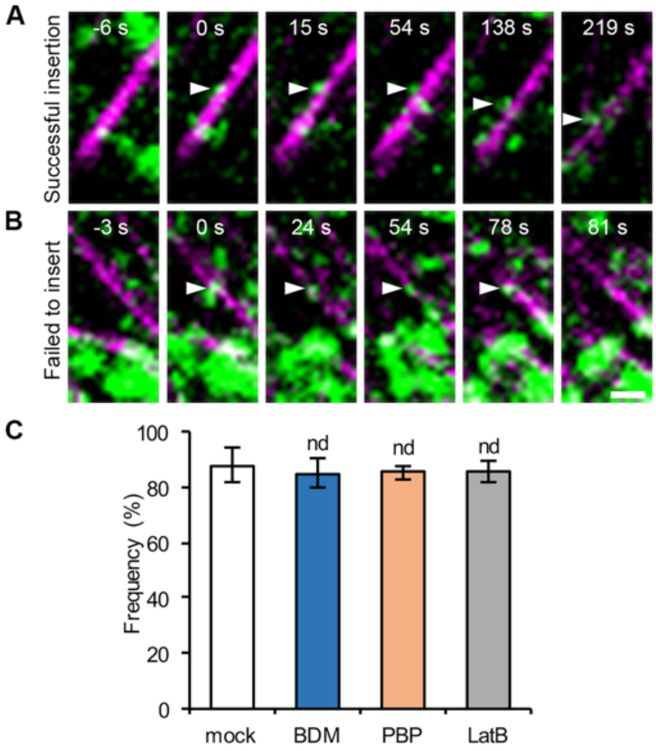


**Supplemental Figure 13.** Inhibition of myosin does not affect the microtubule-dependent positioning of CSC insertion sites.

A and B, Representative images show colocalization of YFP-CESA6 (green) and mCherry-TUA5 (magenta) in etiolated hypocotyl epidermal cells using a co-expression line. A, A successful CSC insertion event shows a newly-arrived CSC particle (white arrowheads) that pauses along a cortical microtubule for ~1 min (0 s, 15 s and 54 s) and then migrates steadily along the microtubule (138 s and 219 s). B, A failed insertion event in inhibitor-treated cells. A CSC particle (white arrowheads) pauses at a cortical microtubule site for more than 1 min (0 s, 24 s, 54 s and 78 s) and then disappears (81 s) without delivering a CSC. Bar = 1 µm. C, Quantitative analysis of the frequency of coincidence of CSC insertion sites with cortical microtubules. The frequency was not altered after BDM, PBP and LatB treatment compared to that in control cells. Values given are means ± SE (n = 5–6 cells per treatment; A total of 53–77 insertion events was measured in each treatment; Student's *t* test, nd: $P > 0.05$).

### Supplemental Movie Legends

**Supplemental Movie 1.** Myosin inhibitor treatments reduce Golgi motility.
Same as Fig. 2A. Time-lapse series were acquired by VAEM at 0.5-s intervals. Video playback rate = 20 frames per second (fps). Total elapsed time = 30 s. Bar = 10 μm.

**Supplemental Movie 2.** Myosin inhibitor treatments reduce myosin XIK motility.
Same as Fig. 2D. Time-lapse series were acquired by SDCM at 80-ms intervals. Video playback rate = 12 fps. Total elapsed time = 3.2 s. Bar = 5 μm.

**Supplemental Movie 3.** Small CESA compartments accumulate in the cortical cytoplasm of myosin- and actin-deficient cells.
Time-lapse series were acquired by SDCM at 0.5-s intervals. CESA compartments in *xi3KO* or inhibitor-treated cells moved rapidly in an erratic or diffusive pattern. Video playback rate = 4 fps. Total elapsed time = 10 s. Bar = 5 μm.

**Supplemental Movie 4.** A successful CSC insertion event at the PM.
Same as Fig. 7A. The CSC particle was marked by yellow arrowheads. Time-lapse series were acquired by SDCM at 3-s intervals. Video playback rate = 20 fps. Total elapsed time = 327 s. Bar = 1 μm.

**Supplemental Movie 5.** A CSC particle is inserted next to a cortical microtubule and then migrates along the microtubule.
Same as Supplemental Fig. S13A. The CSC particle was marked by yellow and white arrowheads. Time-lapse series were acquired by SDCM at 3-s intervals. Video playback rate = 20 fps. Total elapsed time = 237 s. Bar = 2 μm.

**Supplemental Movie 6.** A failed CSC insertion event that had a short pausing time at the PM.
A newly arrived CESA compartment (yellow arrowhead) paused at a fixed location for 30 s and then rapidly moved away. Time-lapse series were acquired by SDCM at 3-s intervals. Video playback rate = 4 fps. Total elapsed time = 72 s. Bar = 2 μm.

**Supplemental Movie 7.** A failed CSC insertion event that had a long pausing time at the PM.
A newly arrived CESA compartment (yellow arrowhead) paused at a fixed location for 210 s and then rapidly moved away. Time-lapse series were acquired by SDCM at 3-s intervals. Video playback rate = 20 fps. Total elapsed time = 270 s. Bar = 2 μm.
